# Information theory of chemotactic agents using both spatial and temporal gradient-sensing

**DOI:** 10.1101/2023.10.14.562229

**Authors:** Julian Rode, Maja Novak, Benjamin M. Friedrich

## Abstract

Biological cells and small organisms navigate in concentration fields of signaling molecules using two fundamental gradient-sensing strategies: spatial comparison of concentrations measured at different positions on their surface, or temporal comparison of concentrations measured at different locations visited along their motion path. It is believed that size and speed dictate which gradient-sensing strategy cells choose, yet this has never been formally proven. Using information theory, we investigate the optimal gradient-sensing mechanism for an ideal chemotactic agent that combines spatial and temporal comparison. We account for physical limits of chemo-sensation: molecule counting noise at physiological concentrations, and motility noise inevitable at the micro-scale. Our simulation data collapses onto an empirical power-law that predicts an optimal weighting of information as function of motility and sensing noise, demonstrating how spatial comparison becomes more beneficial for agents that are large, slow and less persistent. This refines and quantifies the previous heuristic notion. Our idealized model assuming unlimited information processing capabilities serves as a benchmark for the chemotaxis of biological cells.

---

Biological cells employ two different gradient-sensing strategies for chemotaxis [1–3]: *spatial comparison* (SC), where a cell measures the external concentration of signaling molecules simultaneously at different positions on its surface, enabling it to directly estimate a spatial concentration gradient [4–8], and *temporal comparison* (TC), where a cell moves actively and determines whether the local concentration measured sequentially along its path increases or decreases in time [9–13]. Small bacteria such as *E. coli* employ TC, while larger and slower eukaryotic cells commonly employ SC. Gradient-sensing allows these cells to move up concentration gradients. This chemotaxis enables bacteria to forage for food [9–11], social amoeba to aggregate [5–8], or immune cells to migrate to inflammation sites [4]. Sperm cells of marine invertebrates use a mixed strategy of temporal sampling along chiral paths to find the egg [14]. What is common to these examples is the paramount presence of noise.

In seminal work, Berg and Purcell investigated physical limits of chemosensation in light of sensing and motility noise [15]. Cells sense extracellular concentrations by detecting stochastic binding events of signaling molecules to specialized receptors on their surface, which, even if intracellular signaling were perfect, comprises inevitable molecule counting noise. Thermal noise and active fluctuations during cell migration cause motility noise that randomizes cell orientation, rendering previous estimates of gradient direction uncertain [1, 15]. The Berg-Purcell limit was later refined and fundamental sensing limits for either pure SC or TC derived, see [16, 17] for SC, and [18–21] for TC. It was shown that biological cells operate at the information-theoretic limits of gradient sensing during chemotaxis if concentration gradients are shallow, see [6, 7, 22] for SC, and [12, 13, 23, 24] for TC.

Yet, the basic question, which of the two gradient-sensing strategies, SC or TC, provides more information for cells of a given size, is still open. As a first step towards an answer to this long-standing question, we here consider an ideal agent with unlimited information processing capabilities, building on recent advances in information-theoretically optimal navigation strategies [25–32]. While it would be naïve to assume that single biological cells can perform the same information processing tasks, our ideal case serves as base-line for the chemotaxis of biological cells [13], and can give hints on the evolutionary choice of gradient-sensing strategy.

We address both extremes of the exploitation-exploration trade-off, investigating greedy strategies that either maximize *exploitation*, i.e., use of existing information (maximum-likelihood), or maximize *exploration*, i.e., acquisition of additional information (infotaxis). Infotaxis as pioneered by Vergassola et al. [25] harnesses the concept of information maximization from engineering [33], and provides an efficient and reliable heuristic for source-tracking problems [31]. It describes an idealized chemotactic agent that continuously updates a spatial likelihood map of putative target position based on noisy concentration measurements, moving at each time step in the direction that maximizes the expected information gain. We build on a recent extension of infotaxis to spatially extended agents subject to motility noise developed in our group [34], which provides a minimal model of agents performing both SC and TC. Linearization of the originally nonlinear model enables efficient simulations. This allows to dissect the optimal weighting of SC and TC in agents capable of both, akin to a nonlinear Kalman filter that optimally weights different noise-corrupted information sources [35], using as reference scenario static concentration fields established by a single source. We quantitatively show how SC becomes increasingly beneficial for agents of larger size that move slowly and with less directional persistence, suggesting why crawling eukaryotic cells use SC and swimming bacteria use TC.

## Spatial and temporal comparison

Principally, chemotactic agents are characterized by their size *a*, speed *v*, as well as a sensing rate *J*_0_ at which chemoattractant molecules are detected, which may be assumed linear in concentration *c* as *J*_0_ = *λc* [15], with rate constant *λ* that gauges sensing noise. Motility noise randomizes the direction of motion with effective rotational diffusion coefficient *D*_rot_.

An agent of finite size *a* can detect concentration differences across its diameter, with corresponding signal-to-noise ratio of gradient sensing by SC [7, 16, 36]

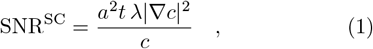

where *t* denotes measurement time.

Alternatively, for gradient-sensing by TC, active motion enables agents to detect concentration changes at a rate *v* ∇*c* along their trajectory, with a signal-to-noise ratio that increases with the cubic power of measurement time *t* [21]

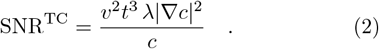

It has been proposed that effective measurement times are set by the rotational diffusion time 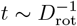 on which swimming direction becomes randomized [1, 37], yet recent works suggest more general power-laws [38–40].

## Bayesian chemotaxis combing spatial and temporal comparison

We present a minimal model of an ideal chemotactic agent combining SC and TC, to dissect the relative importance of SC for finding a target at the center of a radial concentration profile *c*(**x**) of signaling molecules, assuming a baseline scenario of unlimited information processing capabilities. The agent is characterized by three fundamental parameters: size *a*, a rate constant of molecule detection *λ*, and rotational diffusion coefficient *D*_rot_, see Fig. 1 (all other parameters such as speed can be eliminated by re-scaling space or time).

**FIG. 1.**
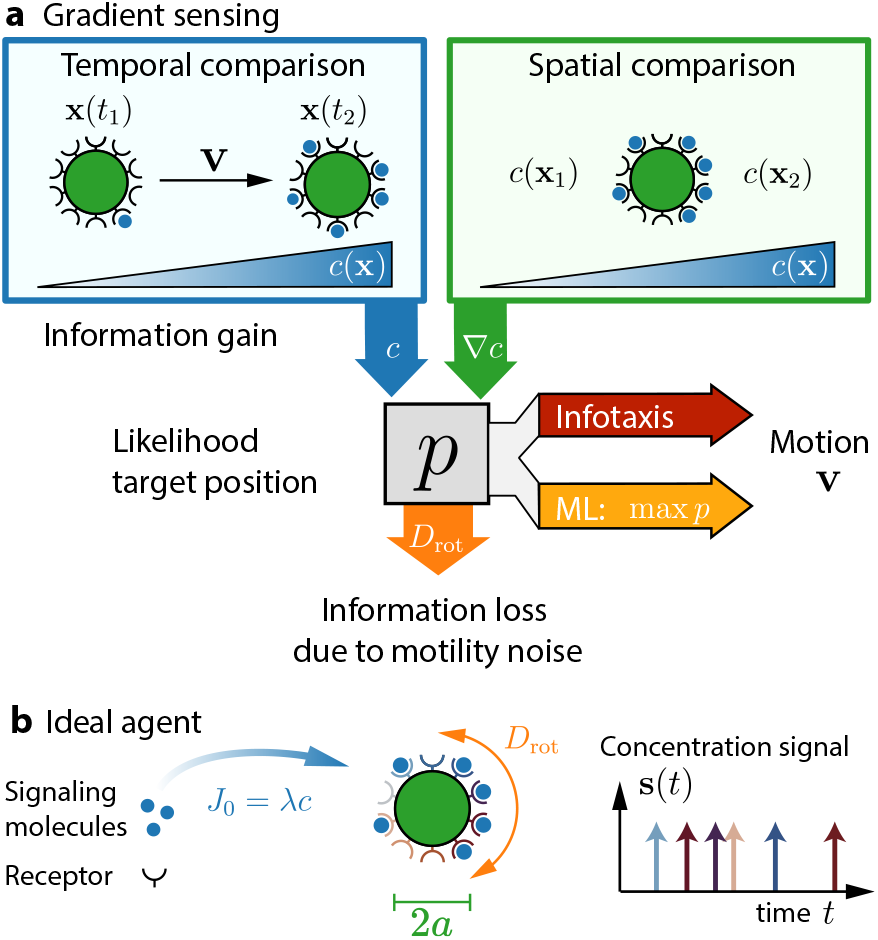
Chemotactic navigation by temporal and spatial comparison. **a** Schematic of gradient-sensing by temporal comparison (TC), where chemotactic agents detect concentration differences along their motion path at different times (left), and spatial comparison (SC), where agents detect concentration differences across their diameter (right). These strategies represent information sources, from which agents can infer a likelihood map *p*(**x**) of target position **x**. Agents take decisions on their next motion step with velocity **v** based on this map using either maximum-likelihood (ML) or infotaxis. Motility noise randomizes motion direction and results in a continuous loss of information. **b** Model of an ideal agent that registers both time and position of signaling molecules binding at rate *λc* to receptors on its surface. The agent has size *a*; motility noise is modeled by an effective rotational diffusion coefficient *D*_rot_.

The agent continuously updates a likelihood map *p*(**x**) of relative target position, based on the timing of molecule detection events *and* the position of detection on its circumference using Bayesian inference, see Eq. (5) in the Method’s section. Different from [25, 27, 29–32], the agent uses a co-moving egocentric map, which allows to study motility noise. The expected change in information of this ideal chemotactic agent is the sum of two terms of information gain for SC and TC, respectively, and a loss term due to motility noise, see Eq. (6). Active motion with velocity **v** changes this expected information gain (to second order in the duration of the time-step), again comprising an SC-term (equal to the spatial gradient of a rate of information gain from SC), and a TC-term (equal to a weighted gradient of the local rate of molecule detection), see (7). This analytical theory, developed in [34], was rewritten here in more compact form to highlight information gain from SC and TC. While the originally nonlinear theory was numerically challenging, a simple linearization proposed here speeds up simulations substantially. Without loss of generality, we assume that agents move with constant speed *v* = |**v**|. Changing *v* can be compensated by rescaling time, which rescales *λ* and *D*_rot_; like-wise, changing concentration *c* is equivalent to changing *λ*.

The agent needs to choose its direction of motion in each time step. We tested two prototypical decision rules, which maximize either exploitation or exploration of information, respectively. In a maximum-likelihood (ML) strategy maximizing exploitation, the agent moves straight towards the global maximum of the current likelihood map *p*(**x**, *t*). Infotaxis provides a second possible navigation strategy, in which the agent chooses its velocity vector **v** such that the expected change in information is maximized [25]. For our spatially extended agent, this is the case if **v** maximizes Eq. (7), which represents two possibly conflicting decision incentives, represented by the SC-term and the TC-term, respectively (see above). In the following, we study both strategies for different noise regimes and agent sizes.

## Stereotypic navigation in absence of motility noise

We first simulated chemotactic agents without motility noise using either the maximum-likelihood strategy or infotaxis, see Fig. 2a. Agents using infotaxis display stereotypic behavior and first move along a straight path whose direction is determined by initial conditions. After a characteristic turning event that defines a time *t*_turn_, these agents home in to the target along approximately circular arcs. In contrast, agents using a maximum-likelihood strategy display more erratic motion before they move approximately ballistically to the target. Snapshots of likelihood maps shown as insets indicate that agents can swiftly estimate target distance, but less so target direction, as reflected by the initially annulus-shaped likelihood map. As agents accumulate directional information, these annulus-shaped distributions become crescent-shaped. Occasionally, distributions become bimodal when agents move towards the center of the crescent (where likelihood is high), but do not sense the expected increase in molecule detection frequency, thus erasing likelihood there, which splits the crescent in two.

**FIG. 2.**
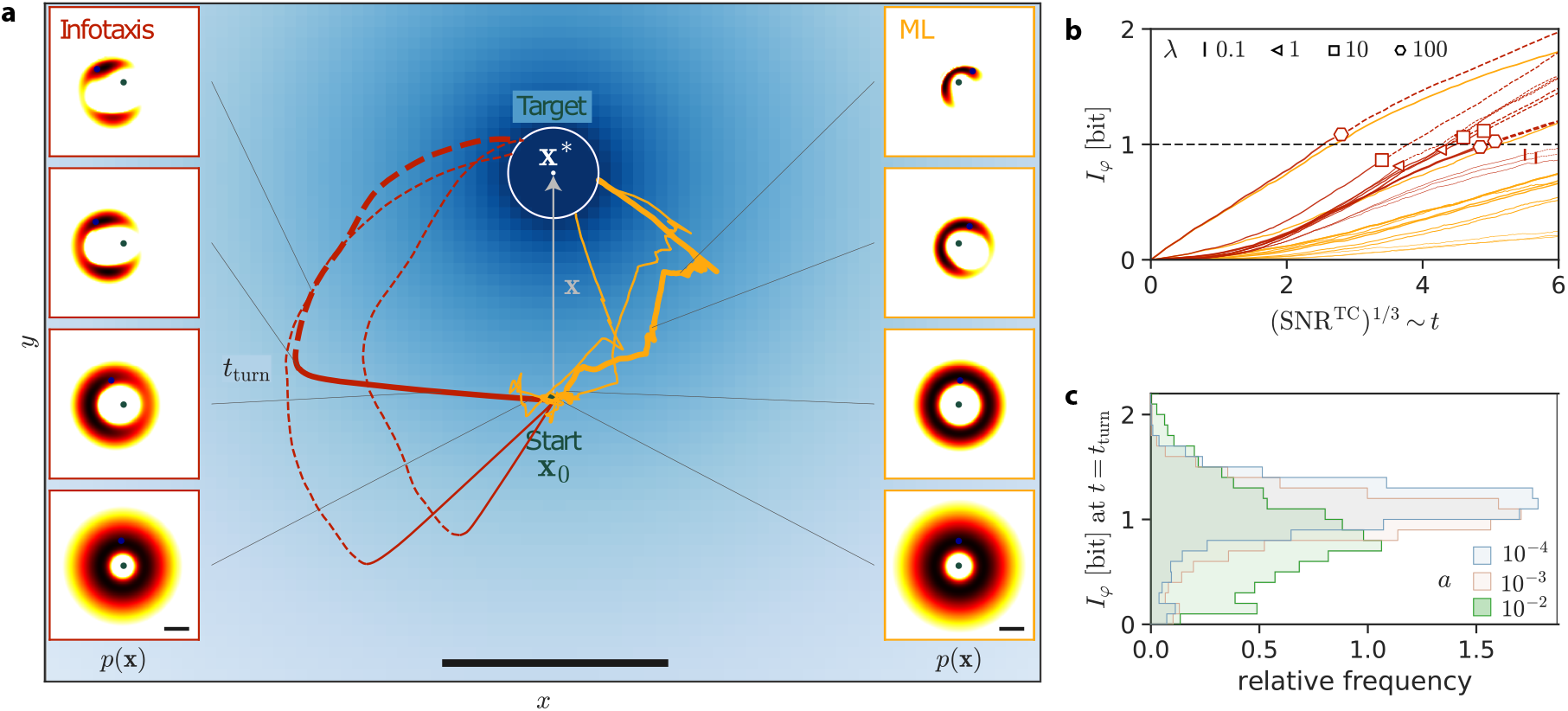
Stereotypic motion without motility noise. **a** Typical trajectories of chemotactic agents (start point: green) navigating in a two-dimensional concentration field *c*(*x, y*) of signaling molecules (blue) released by a target (dark-blue). Agents use either a maximum-likelihood decision rule (ML, orange), or infotaxis maximizing the expected information gain (red). Insets depict likelihood maps *p*(**x**) at selected times for one example trajectory each. For infotaxis, we distinguish a first straight section of the trajectories (solid), and a second section (dashed) after a turning point at time *t*_turn_. Scale bar: 0.2 length units. **b** Directional information *I*_*φ*_ on target direction as function of renormalized time expressed in terms of a signal-to-noise ratio SNR^TC^ of temporal comparison, Eq. (2), for infotaxis (red) and ML (orange), for different values of the rate constant *λ* of molecule detection and agent size *a* (ensemble mean; symbols for infotaxis indicate turning point with symbol shape indicating *λ* according to legend and symbol size increasing with *a*). **c** Distribution of *I*_*φ*_(*t*_turn_) at the first turning point for the case of infotaxis for different agent sizes (represented by line style, see legend; data pooled for different values of *λ*). Parameters: *D*_rot_ = 0, a: *a* = 10^*−*4^, *λ* = 1, b, c: *a* = 10^*−*4^ *−* 10^*−*2^, *λ* = 1 *−* 100.

Bayesian updating is performed on the two-dimensional map of possible target positions, comprising information on both target distance and target direction. The estimated target direction reflects cumulative gradient-sensing by combined TC and SC. To monitor the gain in this directional information, we compute the negative Shannon entropy in bits *I*_*φ*_ = ∫ *dφ p*(*φ*) log_2_[2*π p*(*φ*)], where *p*(*φ*) is the marginal distribution of the polar angle *φ* of relative target direction, see Fig. 2b. For the uniform distribution *p*(*φ*) = 1*/*(2*π*), *I*_*φ*_ = 0, reflecting the complete lack of directional information. When the agent gains information on target direction, and thus *p*(*φ*) becomes more peaked, *I*_*φ*_ increases to become more positive. Indeed, the ensemble average of *I*_*φ*_ increases monotonically as function of time, here normalized using the signal-to-noise ratio SNR^TC^ of TC, Eq. (2). The small spread of curves for infotaxis agents indicates that information gain before the first turning event is dominated by TC.

The characteristic turning events observed for infotaxis agents consistently occur when *I*_*φ*_ reaches the critical value of 1 bit, see Fig. 2c (Fig. S1 for more examples). This is exactly the amount of directional information needed to judge whether the target lies in one half-space or the other.

## Information flow balance with motility noise

Motility noise, modeled as effective rotational diffusion with diffusion coefficient *D*_rot_, sets a finite persistence length *l*_*p*_ = *v/D*_rot_ of agents, see Fig. 3, where *l*_*p*_ = 1 in Fig. 3a and *l*_*p*_ = 0.2 in Fig. 3b. The randomization of swimming direction causes a continuous loss of information on putative target position. The result is a quasi-steady state of information flow, where information gain due to chemosensation and information loss due to motility noise balance each other, reflected by a saturation of directional information *I*_*φ*_(*t*) at a maximal value 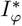, see Fig. 3c, d.

**FIG. 3.**
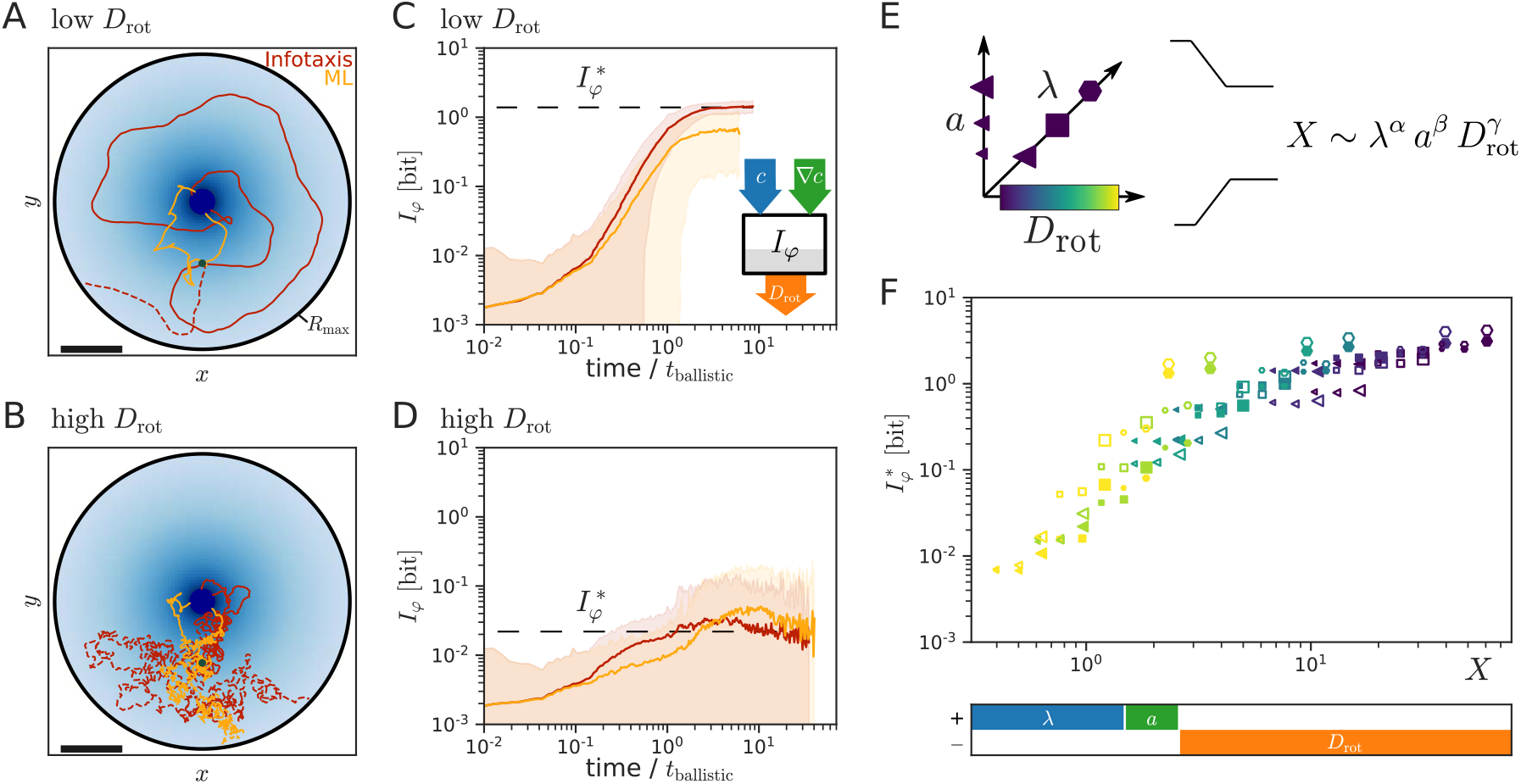
Quasi-steady state of information flow with motility noise. **a, b** Typical trajectories of agents subject to motility noise using either a maximum-likelihood strategy (orange) or infotaxis (red) with low *D*_rot_ (panel **a**, *l*_*p*_ = 1) or high *D*_rot_ (panel **b**, *l*_*p*_ = 0.2). Dashed trajectories failed to find the target, but reached the absorbing boundary (black) at *R*_max_ before. Scale bar: 0.2 length units. **c, d** Directional information *I*_*φ*_ as function of normalized time *t/t*_ballistic_ with *t*_ballistic_ = *R*_target_*/v* for cases of low motility noise (panel **c**) and high motility noise (panel **d**) reveal a quasi-steady state of directional information with saturating directional information 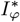 (orange: maximum-likelihood strategy, red: infotaxis; ensemble mean*±* s.d.). The inset in panel **c** sketches information flow determining *I*_*φ*_, with information gained by TC (blue) and SC (green) and lost due to motility noise (orange). **e** The three fundamental model parameters (*λ, D*_rot_, *a*) can be combined into a single empirical power-law 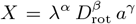 for any simulated quantity such as 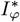 to test if that quantity collapses on a master-curve as function of this single effective parameter *X* (with exponents determined by a linear fit of logarithms). **f** Quasi-steady states of directional information collapse on a master-curve 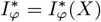 with effective master parameter *X* that obeys an empirical power-law 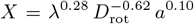 (maximum-likelihood strategy: open symbols, infotaxis: filled symbols; agent parameters encoded according to coordinate system in panel **e**). Power-law exponents were determined for infotaxis agents by a multi-variate fit and are visualized below the figure. Parameters (unless stated otherwise): a, c: *a* = 0.01, *λ* = 1, *D*_rot_ = 0.01; b, d: same as a but *D*_rot_ = 0.5; e, f: *a* = 10^*−*4^ *−* 10^*−*2^, *λ* = 1 *−* 100, *D*_rot_ = 5 *·* 10^*−*3^ *−* 1.

The mean angular information 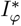 at this quasi-steady state collapses onto a master-curve 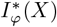 with a master parameter 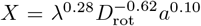 that combines the rate constant *λ* of molecule detection, *D*_rot_, and agent size *a* into an effective power-law, see Fig. 3e, f. Power-law exponents were determined by linear regression of the logarithm of 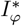 (for infotaxis) as function of the logarithms of the independent variables, which provides a robust collapse on a master-curve even for this nonlinear functional relationship. The behavior for the maximum-likelihood strategy is remarkably similar. The exponents of the power-law reflect the competition between information gain due to TC (characterized by *λ*) and SC (characterized by *λa*), and information loss (characterized by *D*_rot_). A similar power-law holds for the probability of agents to eventually find the target, see Fig. S2 in the Supplemental Material (SM).

## Relative importance of spatial comparison

To gain further insight into the relative importance of temporal comparison (TC) versus spatial comparison (SC) for target finding, we computed their respective contributions to the directional information *I*_*φ*_. While information decomposition remains challenging in general, we can make use of our analytical theory. For each information source, we computed the corresponding information gains 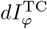 and 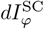 to *I*_*φ*_ using partial updates that involve only that term on the right-hand side of the update equation Eq. (5) that corresponds to this information source. By integrating over trajectories and averaging, we thus obtain the mean cumulated information gains, 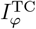 and 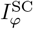, for TC and SC, respectively (conditioned on agents that found the target), see SM text for details. Fig. 4a shows the relative fraction 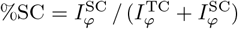 of the information gain for SC as function %SC(*Y*) of a second master parameter obeying a second empirical power-law 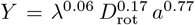 obtained by a linear fit. All three exponents in this power-law are positive. Agent size *a* has the largest exponent and is thus the presiding factor that determines the relative importance of spatial comparison. This is consistent with the strong dependence of the signal-to-noise ratio SNR^SC^ of SC on agent size *a*, see Eq. (1). An increase in motility noise (higher *D*_rot_) like-wise increases the contribution of SC. While rotational diffusion invalidates previously obtained directional information, irrespective of whether it was inferred from SC or TC, TC additionally requires a motion path as straight as possible to probe the concentration field, and is thus affected by reduced directional persistence more strongly. An increase in sensing noise (lower *λ*) slightly reduces the contribution of SC. This indicates a higher vulnerability of SC to sensing noise, consistent with the notion that small concentration differences need to be compared across a relatively small distance in SC.

**FIG. 4.**
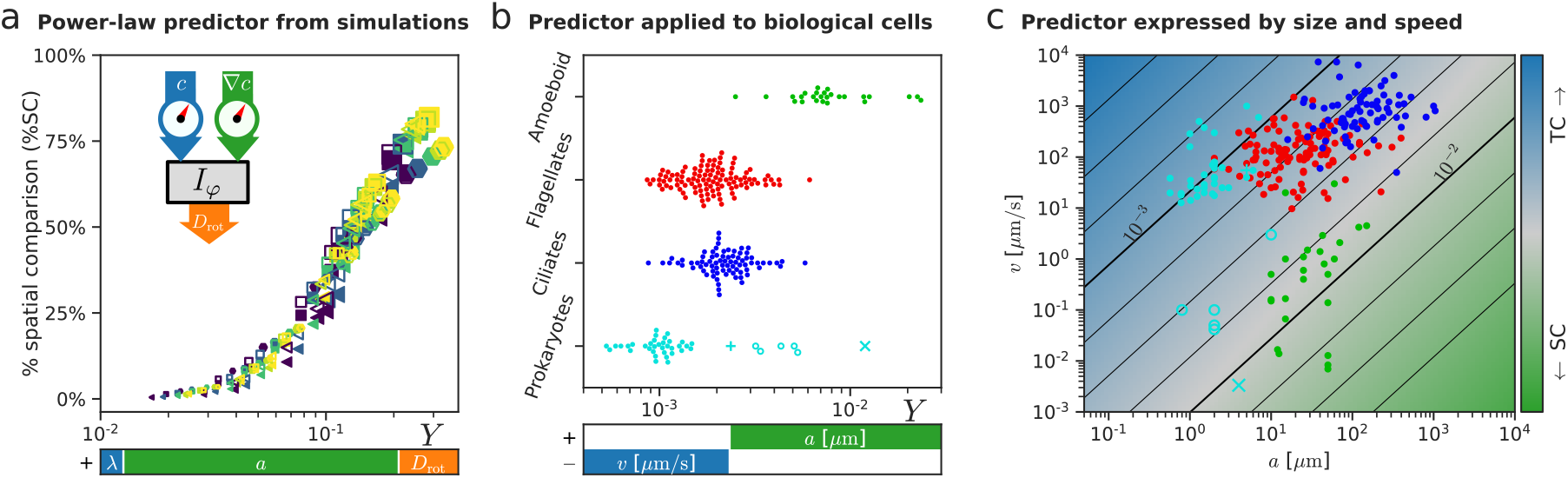
Information decomposition reveals relative importance of spatial comparison. **a** The relative importance of spatial comparison %SC for target finding, computed by metering information flow, collapses on a master-curve %SC(Y) as function of an effective master parameter *Y* that obeys an empirical power-law 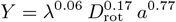, which combines all model parameters (maximum-likelihood strategy: open symbols, infotaxis: filled symbols; agent parameters encoded according to inset in Fig. 3). Specifically, we computed the cumulated gain in directional information using partial update steps corresponding to TC and SC according to Eq. (5), respectively, see text for details. Power-law exponents for *Y* were determined for infotaxis agents by a multi-variate fit and are visualized below the figure. **b** Values of the predictor *Y* estimated for biological cells performing chemotaxis with data taken from [2], using the power-law from panel **a** and making the conservative assumption that rotational diffusion is set by thermal fluctuations. The color code represents different navigation strategies: cyan: bacteria which mostly perform gradient-sensing by temporal comparison (dots), and likely exceptions (other symbols, see text), blue and red: ciliates and flagellates performing helical klinotaxis representing a mixed strategy, green: eukaryotic cells with amoeboid motility performing chemotaxis by spatial comparison. The power-law can be equivalently expressed as *a*^0.32^*/v*^0.23^ in terms of directly measurable parameters size *a* and speed *v*, using an assumption on the diffusion coefficient (see SM). **c** Data from [2] replotted in *a*-*v* parameter space, together with the empirical power-law for the relative importance of spatial comparison as in panel **a** plotted as color code. Parameters: *a* = 10^*−*3^ 2 10^*−*2^, *λ* = 1 100, *D*_rot_ = 0.1 1, panels b, c: reference concentration *c* = 1 nM.

Because our theoretical analysis considers an ideal agent with unlimited information processing capabilities, it is not directly clear whether the behavior of biological cells is dictated by similar laws. Therefore, we reanalyzed data from 252 different cell types corresponding to four groups (prokaryotes, ciliates, flagellates, cells with amoeboid motility) as reviewed in [2], as well as two unusual bacteria described in [41, 42]. Fig. 4b shows for each cell type the value of the power-law derived in Fig. 4a for the ideal agent. Prokaryotes commonly perform gradient-sensing by TC (with exceptions discussed below). Ciliates and flagellates employ a mixed strategy known as helical klinotaxis combining aspects of TC and SC also reviewed below. Cells with amoeboid motility use chemotaxis by SC [5–7].

SC was recently demonstrated for pili-based twitching chemotaxis of the bacterium *Pseudomonas aeruginosa* [41], marked by a star in Fig. 4b. A bacterium described in [42] and marked by a cross in Fig. 4b, displays chemotaxis by SC at very high oxygen concentrations, corresponding to a high effective value of *λ*. Other bacteria that seem to deviate from the rule (open circles in Fig. 4b), likewise display unusual types of motility, such as pili-based motility or actin polymerization, or perform phototaxis instead of chemotaxis [2]. Fig. 4b thus demonstrates that different cells employ temporal or spatial comparison exactly when our idealized theory indicates a higher relative importance of one versus the other.

For the above comparison, agent sizes and speeds as reported in [2, 41, 42] were converted to values of the parameters *λ, D*_rot_, *a*, using a conservative estimate for *D*_rot_ that assumes rotational diffusion is set by thermal fluctuations, hence *D*_rot_ ∼ *a*^*−*3^ [43], while the diffusive flux of signaling molecules to the agent scales linearly with its size, *λ* ∼ *a* [15]. With these assumptions, and a rescaling of time to account for different speeds of cells, we can re-map parameters and re-state the empirical power-law from Fig. 4a in terms of the directly measurable parameters cell size *a* (measured in *μ*m), and speed *v* (measured in *μ*m*/*s), see Fig. 4b bottom and the equivalent representation in Fig. 4c. Explicitly, we used the initial target distance *R*_target_ as typical length-scale and *t*_ballistic_ = *R*_target_*/v* as typical time-scale, to write the power-law from Fig. 4a in dimensionless form

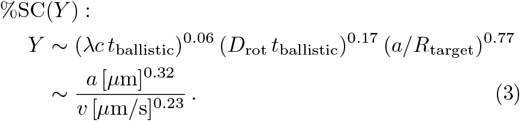

In the second line, we used the scaling of *D*_rot_ and *λ*, as function of the directly measurable quantities size *a* and speed *v*, see SM for details.

By Eq. (3), additional factors can affect the relative importance of SC. The example of a bacterium using SC described in [42] from above is characterized by a 5000-fold increased value of *λc* relative to the reference value used otherwise. Accounting for this high value of *λc* and thus reduced sensing noise reflected in our predictor *Y* separates this unusual bacterium from other prokaryotes that perform TC. If only size *a* and speed *v* are considered as in Fig. 4c, this is not possible.

Ciliates and flagellates shown in Fig. 4b, c employ a navigation strategy known as helical klinotaxis [2, 14, 44, 45]. As in TC with a single effective sensor, these cells sample a scalar signal along their swimming path. Yet, swimming paths are helical, so in the course of a single helical turn, this sensor will have probed all directions, just as agents performing SC, thus enabling directed steering. Eq. (1) could be extended to helical klinotaxis if instead of *a* the radius *R* of helical swimming paths is used. Helical klinotaxis is most efficient in three space dimensions, and only possible if the persistence time 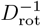 exceeds the period *T* ∼ *R/v* of helical swimming [2, 3].

In conclusion, our information-theoretic analysis explains the previous heuristic observation that large and slow cells commonly employ spatial comparison for gradient-sensing, while small and comparatively fast cells use temporal comparison instead [1, 2, 46].

## A phototactic robot

For demonstrative purposes and as proof-of-principle that bio-inspired Bayesian chemotaxis combining TC and SC may be applied in autonomous search robots (instead of using only TC [47– 49]), we built a phototactic robot. Fig. 5a shows our mecanum-wheeled toy robot capable of motion in all directions, which we equipped with a circular array of 4 light sensors connected to a control unit, see SM for details. Imperfections of the robot drive cause small deviations between control signal and actual movement, which we characterize by an effective rotational diffusion coefficient *D*_rot_, see Fig. S12 in SM. As proxy for a scalar field corrupted by shot noise, the robot navigates in response to a fluctuating pixel pattern projected on the floor, where the flashing probability of each pixel is proportional to *λc*(*x, y*). The robot uses Bayesian inference as in simulations, but neglects its own motility noise as if *D*_rot_ = 0.

**FIG. 5.**
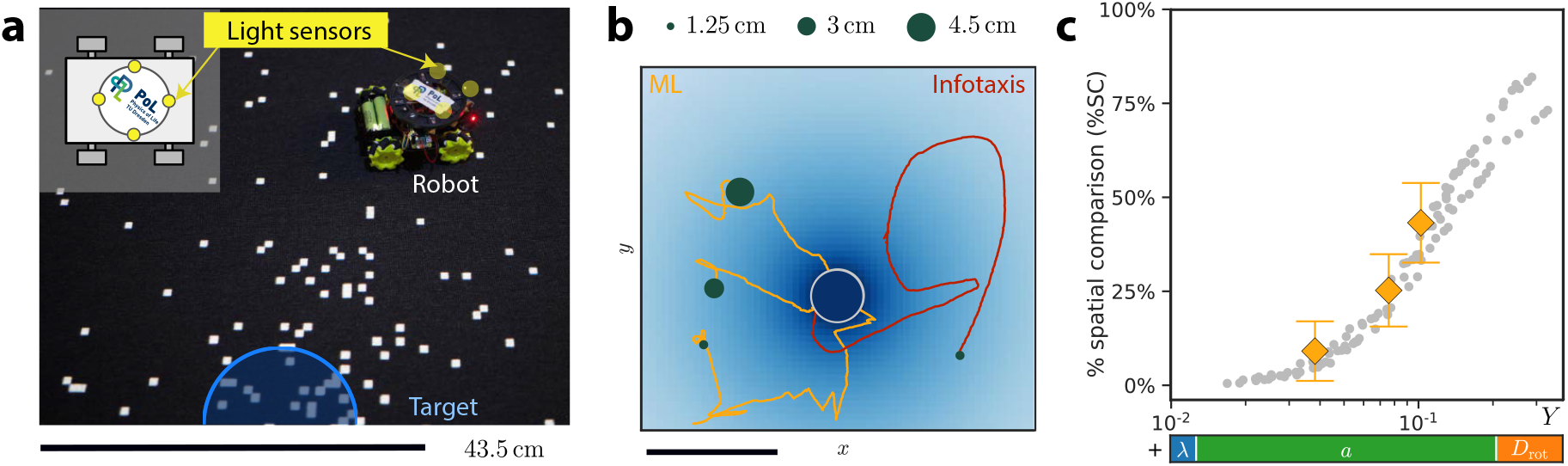
A phototactic robot combining spatial and temporal comparison. **a** Search robot equipped with array of 4 light sensors exposed to a fluctuating pixel pattern projected on the floor. Robots navigate in response to this light stimulus using the same algorithm as ideal agents to find the center of a radial gradient of stochastically flashing pixels. **b** Typical tracks of robots using either maximum-likelihood decision strategy (ML, orange) or infotaxis (red) with different sensor array sizes (green dots mark start positions; tracks rotated for visual clarity). **c** The relative importance of spatial comparison (%SC) for target finding in experiments with ML-robots with different sensor array sizes (orange, mean*±* s.d., *n* = 22, 10, 14) matches simulation data for simulated agents from Fig. 4a (gray dots), using a measured estimate *D*_rot_ = 0.004, see SM for details. Scale bars in a, b: 43.5 cm (which maps to 0.2 length units).

Camera-based tracking yields robot tracks that strongly resemble those in simulations, see Fig. 5b. Changing the size of the sensory array provides a simple means to probe different effective sensing length-scales *a*. The relative importance of spatial comparison in the corresponding robot experiments for different *a* (computed analogously to Fig. 4b), agrees well with simulation results, see Fig. 5c. This demonstrates the feasibility of the approach, despite the complexities of a real robot with actuator noise, which are only partly captured by simulations.

## DISCUSSION

We used information theory to compare two fundamental gradient-sensing strategies for chemotaxis, *spatial comparison* (SC) and *temporal comparison* (TC), by quantifying their relative importance in an ideal agent that combines both. Specifically, a minimal model of an chemotaxis agent with unlimited information processing capabilities allowed us quantify ‘chemotaxis in bits’ in a baseline scenario. An analytical theory allows explicit information decomposition and thus to dissect the relative importance of SC versus TC, which collapses on a master-curve. The corresponding power-law Eq. (3) reflects the expected competition between size *a* and speed *v* [1, 2, 46], additionally tuned by the level of motility noise. The relative importance of SC increases with increasing motility noise, as reduced directional persistence negatively impacts TC.

It could have seem tempting to use the ratio of signal-to-noise ratios SNR^SC^ and SNR^TC^ as a proxy for the relative importance of SC versus TC. However, this ratio depends on the choice of an effective measurement time, which can depend on model parameters in non-trivial ways [38–40]. Our Eq. (3) can thus be interpreted as an empirical power-law for the effective measurement time.

We tested two prototypical decision rules, which represent two extremes of the fundamental exploitation-exploration trade-off: *maximum-likelihood* (ML) and *infotaxis*. Both strategies exhibit similar performance, underscoring the generality of our results. While ML confines agents to the proximity of the target, resulting in a slight overall advantage, infotaxis initially performs better, as the persistent motion of agents right after their start increases information gain from TC. A mixed strategy, which balances exploitation and exploration, may thus optimize performance. Importantly, both strategies yield similar exponents for the empirical power-law on the relative importance of TC versus SC (Table S2 in SM). Similar exponents are also found if a different, exponential concentration field is used (Fig. S6 in SM).

The ideal chemotactic agent introduced here unifies previous theories of either point-like chemotactic agents [20, 21, 25, 27, 31, 32] (only TC), or extended, but mostly stationary agents [16, 17, 39, 50, 51] (only SC). The increased usefulness of SC for larger agents was recently demonstrated in chemotactic agents equipped with small neuronal networks with SC and TC input trained with deep reinforcement learning [52]. An early computational work that addressed the importance of SC versus TC in a one-dimensional model (neglecting motility noise and assuming specific signaling circuits), identified the ratio between cell size and speed already as key discriminator for the relative importance of SC [46], inline with the heuristic rule used in [2] and in agreement with our more general result.

Our analysis thus provides a theoretical underpinning for the previous heuristic notion on the evolutionary choice of different chemotaxis strategies in different cell types [1–3], which could be summarized as small and fast cells use TC, while large and slow (and less persistent) cells use SC. The power-law derived here for a minimal model provides a robust predictor for this choice of chemotaxis strategy in single-celled organisms, see Fig. 4, despite the fact that biological cells have limited information processing capabilities and often need to navigate dynamic gradients [26, 28, 53, 54]. While the optimal weighting of TC and SC could be different for chemotactic agents with limited information processing capabilities, our analysis for ideal agents indicates noise regimes, where SC or TC would provide little additional information, in semi-quantitative agreement with data for biological cells. Moreover, the run-and-tumble motion of the bacterium *E. coli* shares features with the behavior of infotaxis agents in the limit of low motility noise, which comprises a straight run followed by a turn, which would be repeated if there were an information reset after each turn. Biased random walks as observed for amoeboid cells in shallow gradients [6], emerge naturally in our minimal model.

Bayesian chemotaxis requires agents to have a model of typical search environments, yet should be robust with respect to deviations between internal model and reality [39, 55]. Supplemental simulations show that chemotactic agents still exhibit similar performance even if the assumed sensing-noise parameter or rotational diffusion coefficient used in their update step differ from the true values used to generate the input signal and simulate their motion (Figs. S7 and S8 in SM), or if concentration fields are distorted by shear or drift (Figs. S9 and S10). For biological cells, concentration fields may vastly differ for different cells and habitats [26, 53], making it difficult to draw general conclusions. We thus resorted to an idealized, yet prototypical case of a static concentration field shaped by a single source, which allows to compare different chemotaxis strategies in a single model. In biological cells, knowledge about the environment (including characteristic dynamic changes [8, 21, 29, 30, 38, 53, 56]) is likely hard-wired in biochemical signaling networks and learned through evolutionary adaptation. Future work should address the impact of different gradient geometries and the time-scale on which dynamic concentration fields change, yet this is beyond the scope of the current work.

Finally, there exists an extended literature on odor-tracking using sequential Bayesian updating, formulated as partially observable Markov decision processes [29– 33, 57]. This literature uses an equivalent terminology, which we briefly review in the SM. Due to the lack of computationally efficient algorithms, heuristics were proposed to find near-optimal policies [25, 32, 33], of which infotaxis turned out to be surprisingly reliable, efficient, and safe [31]. Other algorithms performed slightly better in specific scenarios, but usually only with a 10%-decrease in mean-first passage time to find a target, and at the expense of reduced robustness [31, 32]. This motivates our use of infotaxis as one of two decision rules tested.

In conclusion, information-theoretically optimal chemotaxis as studied here for an ideal chemotactic agent, which weights gradient-sensing in space and time, provides a baseline scenario to discuss the evolutionary choice of distinct gradient-sensing strategies under the pressure of sensing and motility noise.

Appendix: Analytical theory

We review a minimal model of an ideal chemotactic agent subject to tunable sensing and motility noise previously developed in our group [34], and adapted for efficient simulations here. We first recall the mathematical theory from [34]. Eqs. (5-7) below are directly based on results derived there, though rewritten in more compact form that highlights information gain from SC and TC.

Following [34], we consider a disk-shaped agent of radius *a* and center position x_0_(*t*), which searches for a hidden target at unknown position x^*\**^ = x_0_ + x in the plane. The agent can detect signaling molecules diffusing from the target at each point **x**_**e**_ = **x**_0_ + *a***e** of its circumference at a rate density *λc*(**x**_**e**_)*/*(2*πa*) proportional to the local concentration *c*(**x**_**e**_), where *c*(**x**) ∼ 1*/*| **x** | denotes the steady-state profile established by diffusion from a spherical source. We introduce a vector-valued *chemotactic signal* [39]

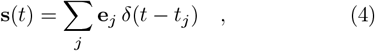

which records binding events occurring at time *t*_*j*_ at position x_0_ + *a* e_*j*_ on the agent’s circumference, modeled as an inhomogeneous Poisson process, see Supplemental Material (SM) for details, and Fig. 1b for visualization. The absolute value *s*(*t*) = |s(*t*) |= ∑ *δ*(*t* − *t*_*j*_) is simply the train of binding events, ignoring their point of detection, as considered in [25]. If the target is assumed to be located at relative position x, the conditional expectation value 𝔼(s | x) = *λa* ∇ *c/*2 of the vector signal s(*t*) encodes the local concentration gradient, while *J* = 𝔼(*s* |x) equals the rate of molecular binding events. This rate is proportional to the mean concentration along the circumference of the agent, *J* ≈ *λ*[*c*(x_0_) + (*a*^2^*/*4) ∇^2^*c*(x_0_)] [34].

The agent moves inside the concentration field with constant speed *v* = |v|, subject to rotational diffusion with effective rotational diffusion coefficient *D*_rot_, providing a minimal model of motility noise [39]. This dynamics extends the popular model of an Active Brownian Particle (ABP) [58], as our agent shall continuously decide on its direction of motion based on the directional chemo-sensory input it receives.

We are interested in the information-theoretic limits of chemotaxis, and therefore assume unlimited information processing capabilities of the agent, which can store and update a likelihood map *p*(x, *t*) of relative target position x = x^*\**^−x_0_. The choice of an *egocentric map*, in which the moving agent is always at the center (with coordinate axes aligned to a co-moving material frame of the agent), is mathematically convenient. We assume that the agent has implicit knowledge of stereotypic concentration fields, and hence knows the conditional rate *J* (x_0_ | x^*\**^) and concentration gradient 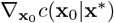, provided the target were at x^*\**^ = x_0_ + x. Here, *c* points towards the target.

Bayes’ rule rationalizes how to update previous beliefs in light of new information. In a time-continuous formulation, the time-evolution of a likelihood map *p*(**x**, *t*) of relative target position reads

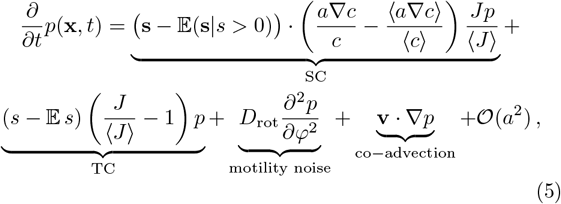

which follows from [34] after linearization by moment-closure, see SM for details. This approximation is key for efficient numerical simulations. Different from [34] (and different from the usual Bayesian update scheme), we can now integrate the dynamics using substantially longer time-steps that include multiple binding events. We confirmed in initial simulations that the full nonlinear scheme and our linearization gave essentially identical results. The expectation values 𝔼 *s* and 𝔼(s | *s >* 0) denote measurements expected by the agent given its current likelihood map *p*(x, *t*), while ⟨·⟩ averages over x, again using *p*(x, *t*). Thus, 𝔼 *s* =⟨ 𝔼(*s* | x) ⟩. Each summand on the right-hand side of Eq. (5) has a straight-forward physical interpretation. The third summand describes loss of information due to motility noise characterized as rotational diffusion, where *φ* denotes the polar angle of x. The last summand describes the motion-induced shift of the egocentric map, in which the the relative position of the target changes as 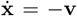 when the agent moves with velocity v. The TC-term describes the update of target position based on the detection of signaling molecules, yet without reference to their position of detection on the circumference of the agent, thus generalizing TC. This term is equivalent to an analytical theory developed previously in [27] for the point-like agents introduced in [25], and has a simple geometric interpretation: whenever binding events are recorded at a rate higher than the expected detection rate (i.e., *s>* ⟨*J* ⟩), the likelihood *p*(x) is increased for all target positions x, for which the local detection rate is higher than this mean rate (*J* (x)*>* ⟨*J* ⟩), but decreased for all other positions. For *s<* ⟨*J* ⟩, this update has the opposite sign. In particular, even the absence of a binding event provides information to the agent, namely that the target might be further away than expected [59].

The SC-term is novel, and describes the update due to spatial comparison of concentration differences across the diameter of the agent. This term is a scalar product of an innovation vector, formed by the difference of the vector-valued concentration measurement **s**(*t*) and its conditional expectation value E(s| *s >* 0) = Es *s/* ⟨*J* ⟩ if a binding event has happened, and the local concentration gradient, likewise corrected by a suitable expectation baseline. If no binding event is detected, the innovation vector is zero, reflecting the fact that it is impossible to record the position of a binding event without noticing that there has been a binding event in the first place.

## Expected change in information

From the current prior *p*(x, *t*), the agent can infer the likelihood to detect a signaling molecule at a particular position, and thus forecast how the negative Shannon entropy *I* = ∫*d***x** *p*(**x**) ln *p*(**x**) is expected to change in the infinitesimal time interval [*t, t* + *dt*] ahead, see [34] and SM text for details

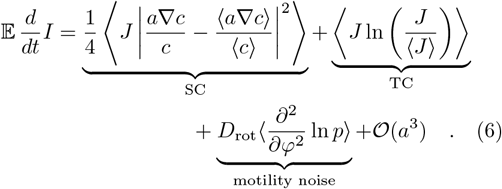

The SC-term characterizes information gain due to spatial comparison and is always non-negative. If multiplied by a characteristic sensing time, this term generalizes the signal-to-noise ratio SNR^SC^ of spatial comparison, see Eq. (1), by accounting for the expected gradient base-line ⟨∇*c*⟩. The TC-term characterizes information gain due to temporal comparison; it is always ≥ 0 because it is proportional to the Kullback-Leibler divergence between the likelihood maps *p* and *p J/* ⟨*J* ⟩ before and after a binding event, respectively. The last summand characterizes information loss due to motility noise; this Fisher information is always ≤ 0 by Bruijn’s inequality [34].

Active motility with velocity v affects the expected change of information only to second order in the time-step, which can be expressed in terms of the time derivative of the expected change in information

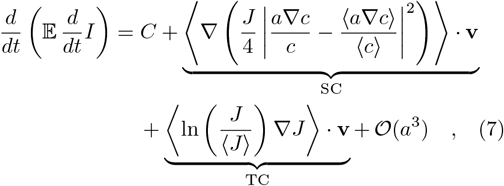

where *C* combines terms independent of **v**. Eq. (7) reformulates a result in [34] in more compact form, see SM for details. The first term characterizes information gain due to SC, whereas the second term characterizes information gain due to concentration sensing akin to TC.

## Data, Materials and Software Availability

Simulation data and Python code for efficient simulation of Bayesian chemotaxis investigated here is available at https://zenodo.org/record/8421582 under DOI 10.5281/zenodo.8421582. Data analyzed in Fig. 4 on speed and size of chemotactic cells is available as Supplemental Data of [2].

## Supporting information

Supplemental text including 12 supplemental figures

Agent without motility noise performing infotaxis.

Agent without motility noise using maximum-likelihood decision rule.

Agent with motility noise performing infotaxis.

Agent with motility noise using maximum-likelihood decision rule.

Robot experiment for maximum-likelihood strategy.

## Acknowledgment

JR, MN, BMF were supported by the German National Science Foundation (DFG) (grant no. 391963627), the *Center for Advancing Electronics Dresden* (cfaed), and the Excellence Initiative by the German Federal and State Governments (Clusters of Excellence cfaed EXC-1056 and *Physics of Life* EXC-2068-390729961). BMF is additionally supported by the DFG through a Heisenberg grant (421143374). We thank Andrea Auconi for stimulating discussions. We thank Camillo Oeser and Arthur Willert for help with experiments, as well as the *Center for Information Services* at TU Dresden for granting use of their High Performance Computing facilities.

