## Supplemental text including 12 supplemental figures for "Information theory of chemotactic agents using both spatial and temporal gradient-sensing"

#### A. DETAILS ON NUMERICAL SIMULATIONS

*Search domain.* For numerical simulations of agents performing Bayesian chemotaxis, we consider disk-shaped agents of radius $a$ that can move freely in a disk-shaped, two-dimensional search domain with absorbing boundary conditions at a maximal radius $R_{\max}$. A single target is located at the center of the search domain. We assume a radial concentration profile $c(\mathbf{x}) = 1/|\mathbf{x}|$ corresponding to the steady-state profile established by diffusion from a spherical source [45]. To ensure comparability of simulations runs, agents start at a fixed position $\mathbf{x}^* = (0, -R_0)^T$ with distance $R_0 = 0.2$ from the target.

*Egocentric map.* The egocentric likelihood map of the agent likewise has dimensions of a disk with radius $R_{\max}$. This egocentric map has its origin at the midpoint of the agent, with coordinate axes aligned to a material frame of the agent characterized by orthonormal vectors $\mathbf{h}_1$ and $\mathbf{h}_2$. Relative coordinates are expressed as $\mathbf{x} = x_1\,\mathbf{h}_1 + x_2\,\mathbf{h}_2$. The agent does not move in its egocentric map, while the target is translated by $-\mathbf{v}\,\Delta t$ in each time-step $\Delta t$ due to the active motion of the agent, as well as rotated by a random angle $\Delta\varphi$ around the midpoint of the agent to account for the agent's rotational diffusion, where $\Delta\varphi$ is drawn from a normal distribution $\mathcal{N}(0, 2D_{\mathrm{rot}}\,\Delta t)$ with mean 0 and variance $2D_{\mathrm{rot}}$.

*Molecule detection.* The agent has $N_\theta = 50$ detection angles, i.e. receptors with respective directions $\mathbf{e}_j = (\cos\theta_j, \sin\theta_j)^T$ with $\theta_j = 2\pi\,j/N_\theta$. For each receptor, the number of molecular detection events in a time-step $dt$ is drawn from a Poisson distribution with mean $\lambda c(\mathbf{x} - a\,\mathbf{e}_i)\,\Delta t/N_\theta$. Importantly, by using a linearized theory as given in Eq. (5), it is admissible if more than one detection event occurs in a single time-step, which allows to choose substantially larger time-steps $dt$ than in the original nonlinear theory [34], which speeds up simulations substantially.

*Euler scheme for numeric integration.* To integrate the time evolution equation Eq. (5) for the likelihood $p(\mathbf{x}, t)$ of relative target position $\mathbf{x}$, we use a finite difference scheme with forward Euler method. For all simulations, we used a fixed time step of $\Delta t = 0.005$, and spatial step size $\Delta x = \Delta y = 0.01$ (except $\Delta t = 0.0005$ for $\lambda = 100$).

*Rotational diffusion.* The numerically most challenging part is rotational diffusion.

In case of a small rotational diffusion coefficient $D_{\mathrm{rot}} < 0.1$, we use a 9-point stencil for the second angular derivative in Euclidean coordinates

$$\mathcal{D}_{\mathrm{rot}} = \frac{\partial^2}{\partial\varphi^2} = -x_1\partial_{x_1} - x_2\partial_{x_2} + x_2^2\partial_{x_1}^2 + x_1^2\partial_{x_2}^2 - 2x_1x_2\partial_{x_1}\partial_{x_2} \quad . \tag{S1}$$

We note that the pre-factors in front of the second derivatives scale with squared distance $|\mathbf{x}|^2$, which can become as large as $R_{\max}^2$; this can cause numerical problems even for intermediate $D_{\mathrm{rot}}$. For increased numerical accuracy, we therefore implement the rotational diffusion step as $n$ consecutive updates with smaller time-step $dt/n$, where $n$ is the smallest integer that fulfills the Courant criterion $(8\,R_{\max}^2 D_{\mathrm{rot}}\,dt/n)/\Delta x^2 < 0.8$.

In case of larger rotational diffusion coefficients $D_{\mathrm{rot}} > 0.1$, we switch to a kernel-based method for reasons of efficiency. The corresponding rotational diffusion kernel is pre-computed for each point $(x_1, x_2)$ on the simulation grid as

$$K(x_1', x_2'|x_1, x_2) = \int_0^{\Delta t} dt\,\mathcal{D}_{\mathrm{rot}}\,p(x_1', x_2', t|x_1, x_2) \quad , \tag{S2}$$

where initially $p(x_1', x_2', t = 0|x_1, x_2) = \delta(x_1', x_2'|x_1, x_2)$ is zero everywhere except on the grid point $(x_1, x_2)$ where it is one. To reduce memory needs and accelerate the computations, values of $K$ with relative magnitude smaller than $10^{-4}$ were set to zero.

*Initial conditions.* The initial likelihood $p(\mathbf{x}, t = 0)$ is chosen as an almost flat normal distribution (with standard deviation $\sigma \approx 166$). To probe different initial directions, this initial likelihood was shifted by a small distance $\Delta x = 10^{-3}$ along a pre-specified initial direction. We used 7 initial directions with direction angles $\varphi_0 = j\,\pi/4$ with $j \in \{0, 1, 2, 3, 5, 6, 7\}$, where we excluded the initial direction $\varphi = -\pi$ pointing directly towards the target.

*Parameters for Fig. 2.* For simulations without rational diffusion shown in Fig. 2, the absorbing boundary is set at a radius of $R_{\mathrm{max}} = 1.98$, concentric with the target. This large radius ensured that no agent ever reached the absorbing boundary, thus realizing an effectively unbounded simulation domain. We tested $4 \times 3 = 12$ parameter combinations with $\lambda \in \{0.1, 1, 10, 100\}$ and $a \in \{10^{-4}, 10^{-3}, 10^{-2}\}$, with 100 realizations for each of the 7 initial directions angles $\varphi_0$.

*Parameters for Fig. 3.* For simulations with rotational diffusion as shown in Fig. 3, we reduced the radius of the absorbing boundary to $R_{\mathrm{max}} = 0.48$ and increased the number of realizations per initial direction to 200 in order to reduce the computational cost and achieve good statistics. We tested $3 \times 3 \times 5 = 45$ parameter combinations with $\lambda \in \{1, 10, 100\}$, $a \in \{10^{-4}, 10^{-3}, 10^{-2}\}$, and $D_{\mathrm{rot}} \in \{5 \cdot 10^{-3}, 10^{-2}, 5 \cdot 10^{-2}, 10^{-1}, 5 \cdot 10^{-1}, 1\}$ with 200 realizations for each of the 7 initial directions angles $\varphi_0$. We confirmed that choosing a larger radius $R_{\mathrm{max}}$ for the absorbing boundary increases $P_{\mathrm{reached}}$ as expected, but otherwise does not change the behavior of chemotactic agents, see figure S4.

*Parameters for Fig. 4.* As for Fig. 3, we used $R_{\mathrm{max}} = 0.48$ for the radius of the search domain. The range of parameters for Fig. 4 were chosen to cover the cross-over region, where information gain from either SC or TC becomes relevant, see section *Relative importance of spatial comparison* in the main text. Specifically, we tested $3 \times 7 \times 5 = 105$ parameter combinations with $\lambda \in \{1, 10, 100\}$, $a \in \{10^{-3}, 2 \cdot 10^{-3}, 3 \cdot 10^{-3}, 6 \cdot 10^{-3}, 9 \cdot 10^{-3}, 10^{-2}, 2 \cdot 10^{-2}\}$, and $D_{\mathrm{rot}} \in \{0.1, 0.2, 0.5, 0.8, 1\}$ with 20 realizations for each of the 7 initial directions angles $\varphi_0$. From these 140 simulated agents, even for the most unfavorable parameters (high $D_{\mathrm{rot}}$, low $\lambda$), 33 and 115 agents found the target for the infotaxis and maximum-likelihood decision strategy, respectively, ensuring sufficient statistics for conditional expectation values. Simulating 140 realizations for 105 parameter combinations takes about 2000 core hours on the ZIH cluster Barnard (Intel Xeon Platinum 8470).

Default values of parameters used in numerical simulations for Figs. 2-4 are summarized in Table S1.

| Parameter | Default value | Meaning |
|---|---|---|
| $a$ | $10^{-4} - 10^{-2}$, Fig. 4: $10^{-4} - 2 \cdot 10^{-2}$ | agent size (radius) |
| $\lambda$ | $1 - 100$ | rate constant of molecule detection |
| $D_{\mathrm{rot}}$ | Fig. 2: 0, Fig. 3: $5 \cdot 10^{-3} - 1$, Fig. 4: $0.1 - 1$ | rotational diffusion coefficient of agent |
| $v$ | 0.01 | speed of agent |
| $R_{\mathrm{target}}$ | 0.04 | target size (radius) |
| $R_0$ | 0.2 | initial distance of agent from target |
| $R_{\mathrm{max}}$ | Fig. 2: 1.98, Figs. 3,4: 0.48 | radius of circular search domain |
| $\Delta t$ | 0.005 ($\Delta t = 0.0005$ for $\lambda = 100$) | time-step in simulations |
| $\Delta x = \Delta y$ | 0.01 | spatial grid size in simulations |

TABLE S1. Default parameter values used in numerical simulations.

### B. DATA ANALYSIS OF NUMERICAL RESULTS

#### 1. Detection of first turn for agents without motility noise

For agents performing infotaxis without motility noise as shown in Fig. 2, we used the following method to detect their first turn. We first smoothed the trajectories by minimizing the residual between the first derivative of the trajectory and the first derivative of the smooth curve, using the `powersmooth` python package available at `https://github.com/rodjul42/powersmooth`. The regularization weight that determines the strength of smoothing, was chosen as 1000. We compute the curvature of smoothed trajectories as $\kappa = |\dot{x}\ddot{y} - \dot{y}\ddot{x}|/v^3$, and detect peaks of the curvature using a threshold of 15. Occasionally, spurious curvature peaks are detected right at the beginning of trajectories; therefore, peaks followed by further peaks were ignored if the directional information $I_\varphi$ at their time-point is below 0.1.

#### 2. Quantification of information flow

*Directional information.* To compute the directional information $I_\varphi$, we first determined the marginal distribution $p(\varphi) = \int dr\, p(r,\varphi)$ of the polar angle $\varphi$ of the relative target direction $\mathbf{x} = |\mathbf{x}|(\cos\varphi, \sin\varphi)$. For this, we map each pixel $(x_1, x_2)$ of the discrete likelihood map $p(x_1, x_2)$ to its corresponding angular bin $\varphi_{x_1,x+2} \in [\varphi_k, \varphi_k + \Delta\varphi]$ with bin width $\Delta\varphi = \pi/50$, where $\varphi_{x_1,x_2} = \arctan(x_2/x_1)$ is the polar angle of the pixel. This can be efficiently implemented using pre-computed look-up lists of pixel indices. The value of the marginal distribution $p(\varphi_k)$ for each bin is then computed as the sum of the likelihood densities $p(x_1, x_2)$ of the corresponding pixels, multiplied by pixel area $dA$, $p(\varphi_k) = \sum' p(x_1, x_2)\, dA$. Finally, the directional information is computed as the negative Shannon entropy

$$I_\varphi = \mathcal{I}_\varphi[p] = \sum_k p(\varphi_k)\, \log_2\left[2\pi\, p(\varphi_k)\right] \quad . \tag{S3}$$

*Computation of quasi-steady state of directional information $I_\varphi^*$.* To determine the saturation value $I_\varphi^*$ of the time-dependent directional information $I_\varphi(t)$, we fit a clipped linear function $f(t)$ to the ensemble-average of the directional information

$$f(t) = \begin{cases} a\,t & \text{if } t < t^* \\ a\,t^* & \text{else} \end{cases} \quad . \tag{S4}$$

The saturation values is then simply given by $I_\varphi^* = a\,t^*$.

#### 3. Information decomposition

*Relative importance of spatial comparison.* We can compute an information decomposition of the directional information $I_\varphi$ using our analytical theory Eq. (5). We quote Eq. (5) for reference, noting the contribution to the change of $p(\mathbf{x}, t)$ due to rotational diffusion (rot), advection (adv), concentration sensing (TC), and gradient sensing (SC)

$$\frac{\partial}{\partial t} p(\mathbf{x}, t) = \underbrace{D_{\mathrm{rot}} \frac{\partial^2 p}{\partial \varphi^2}}_{\mathrm{rot}} + \underbrace{\mathbf{v}\cdot\nabla p}_{\mathrm{adv}} + \underbrace{p\,(s - \mathbb{E}\, s)\left(\frac{J}{\langle J\rangle} - 1\right)}_{\mathrm{TC}} + \underbrace{\frac{p\,J}{\langle J\rangle}\,(\mathbf{s} - \mathbb{E}(\mathbf{s}|s>0))\cdot\left(\frac{a\nabla c}{c} - \frac{\langle a\nabla c\rangle}{\langle c\rangle}\right)}_{\mathrm{SC}} + \mathcal{O}(a^2) \quad .$$

During each time-step of duration $\Delta t$, we apply the different summands in Eq. (5) sequentially, and track the corresponding changes $\Delta I_\varphi^{\mathrm{rot}}$ due to rotational diffusion, $\Delta I_\varphi^{\mathrm{adv}}$ due to the advection, $\Delta I_\varphi^{\mathrm{TC}}$ due to concentration sensing (TC), and $\Delta I_\varphi^{\mathrm{SC}}$ due to gradient sensing (SC); additionally, we compute the change $\Delta I_\varphi^{\mathrm{TC+SC}}$ for a full update step.

Specifically, we first separate the contributions to the prediction step, (S25), and track the corresponding changes $\Delta I_\varphi^{\mathrm{rot}}$ and $\Delta I_\varphi^{\mathrm{adv}}$ in directional information. Second, we separate the update step, which updates the predicted likelihood $p(\mathbf{x}, t + \Delta t)$ according to the last measurement as in Eqs. (S26) and (S33), into contributions due to TC and SC, and track the corresponding changes $\Delta I_\varphi^{\mathrm{TC}}$ and $\Delta I_\varphi^{\mathrm{SC}}$ in directional information. Thus,

$$\Delta I_\varphi^{\mathrm{rot}} = \mathcal{I}_\varphi\,\underbrace{[p + dt\,\mathcal{D}_{\mathrm{rot}}(p)]}_{p_{\mathcal{D}}} - \mathcal{I}_\varphi[p] \tag{S5}$$

$$\Delta I_\varphi^{\mathrm{adv}} = \mathcal{I}_\varphi\big[\underbrace{p_{\mathcal{D}} + dt\,\mathbf{v}\cdot\nabla p}_{p'}\big] - \mathcal{I}_\varphi[p_{\mathcal{D}}] \tag{S6}$$

$$\Delta I_\varphi^{\mathrm{TC+SC}} = \mathcal{I}_\varphi[p' + \text{full update}] - \mathcal{I}_\varphi[p'] \tag{S7}$$

$$\Delta I_\varphi^{\mathrm{TC}} = \mathcal{I}_\varphi[p' + \text{update TC}] - \mathcal{I}_\varphi[p'] \tag{S8}$$

$$\Delta I_\varphi^{\mathrm{SC}} = \mathcal{I}_\varphi[p' + \text{update SC}] - \mathcal{I}_\varphi[p'] \quad , \tag{S9}$$

where $p = p(\mathbf{x}, t)$ and $p' = p'(\mathbf{x}, t + \Delta t)$ denote the likelihood map before and after the prediction step, respectively, and $\mathcal{I}_\varphi$ denotes the directional information operator, see (S3).

Summing up these respective increments for each time-step yields cumulative information gains for each contribution corresponding to the different terms in Eq. (5). The discrete nature of individual detection events causes jumps in the time-dependent likelihood, which do not scale with the small time-step $\Delta t$. Thus, it is not clear a priori if the sum of the changes in directional information due to either TC or SC equal the corresponding change in directional

| decision rule | exponent for | | |
|---|---|---|---|
| | $\lambda$ | $a$ | $D_{\text{rot}}$ |
| *infotaxis* | 0.06 | 0.77 | 0.17 |
| *maximum-likelihood* | 0.04 | 0.80 | 0.16 |
| *infotaxis – for exponential gradient* | 0.11 | 0.69 | 0.20 |
| *maximum-likelihood – for exponential gradient* | 0.08 | 0.77 | 0.15 |

TABLE S2. Fitted exponents of empirical power-law $Y \sim \lambda^\alpha a^\beta D_{\text{rot}}^\gamma$ for the relative importance %SC of spatial comparison for *infotaxis* agents as shown in Eq. (3) of the main text (first row), as well as analogous exponents for agents using the *maximum-likelihood* decision rule (second row). The last two rows display analogous exponents of the empirical power-law if the exponential concentration profile given by Eq. (S18) is used in the simulations (instead of the concentration field $c(\mathbf{x}) = 1/|\mathbf{x}|$ used in Figs. 2-5). Corresponding master curves for %SC using the exponents of infotaxis agents are shown for the two concentration fields in Fig. 4a and Fig. S6, respectively.

information for the full update step, i.e., whether $\sum \Delta I_\varphi^{\text{TC+SC}} \approx \sum \Delta I_\varphi^{\text{TC}} + \sum \Delta I_\varphi^{\text{SC}}$. Fig. S3 displays cumulated changes in directional information, showing that this decomposition holds true to very good approximation.

We used the ensemble-average of the total cumulated change in directional information, conditioned on agents finding the target, to compute the total information gain due to either TC or SC, $I_\varphi^{\text{TC}} = \overline{\sum \Delta I_\varphi^{\text{TC}}}$ and $I_\varphi^{\text{SC}} = \overline{\sum \Delta I_\varphi^{\text{SC}}}$, from which we determined the relative importance of SC as the ratio of total information gains $I_\varphi^{\text{SC}}/(I_\varphi^{\text{SC}} + I_\varphi^{\text{TC}})$. $I_\varphi^{\text{SC}}/(I_\varphi^{\text{SC}} + I_\varphi^{\text{TC}})$.

We restate the fitted exponents for the empirical power-law for the relative importance of SC for the case of infotaxis as stated in Eq. (3) in the main text, as well as the corresponding exponents for the case of a maximum-likelihood decision rule, see Table S2.

#### 4. Directional information and chemotactic index

The directional information $I_\varphi$ considered here is correlated to the *chemotactic index* CI used by some authors [6, 7, 16], where CI equals the net drift up-gradient in an ensemble of agents normalized by their speed, CI $= \langle \mathbf{v} \cdot \nabla c \rangle / (v|\nabla c|)$. To make this relationship explicit, we restrict to a one-parameter class of prototypical angular distributions, the von-Mises distribution $p(\varphi) \sim \exp[\kappa \cos(\varphi - \varphi_0)]$, where the parameter $\kappa$ describes an inverse dispersion. In fact, the von-Mises distribution is the maximum-entropy distribution for given circular mean $\varphi_0$, and circular variance $\text{var}(\varphi) = 1 - |\oint d\varphi \, \exp[i\,\varphi] p(\varphi)|$; its directional information is given by $[\kappa I_1(\kappa)/I_0(\kappa) - \ln I_0(\kappa)]/\ln(2)$, where $I_0$ and $I_1$ denote modified Bessel functions. If we assume that the likelihood distribution of a chemotactic agent represents a faithful distribution for the direction estimates in an ensemble of agents, we find that a directional information of 0.5 bit and 1 bit, respectively, would correspond to a chemotactic index of approximately 0.56 and 0.75 (with $\kappa \approx 1.37$ and $\kappa \approx 2.40$).

### C. MAPPING PARAMETERS FOR WAN ET AL. [2]

Wan et al. surveys swimming speed and cell size for 252 single-celled organisms [2]. To relate these experimentally directly accessible parameters to the dimensionless parameters in our model, we introduce a constant length-scale $L$, constant concentration-scale $C$, and variable time-scale $T$. This allows us to convert dimensionless model parameters to their dimensionalized counter-parts as follows

$$a \to a\,[\mu\text{m}] = aL \tag{S10}$$

$$v \to v\,[\mu\text{m/s}] = vL/T \tag{S11}$$

$$c \to c\,[\text{nM}] = c\,C \tag{S12}$$

$$\lambda \to \lambda\,[\text{nM}^{-1}\,\text{s}^{-1}] = \lambda/(CT) \tag{S13}$$

$$D_{\text{rot}} \to D_{\text{rot}}\,[\text{s}^{-1}] = D_{\text{rot}}/T \quad . \tag{S14}$$

We choose $L = 12.5 \cdot 10^3\,\mu$m to match the range of cell sizes reported in [2] to the range of agent sizes $a$ used in our simulations, while $T$ is determined for each cell individually to match $v = 0.01$.

Further, we make the following assumptions about the rate constant $\lambda$ of molecule detection and the rotational diffusion coefficient $D_{\mathrm{rot}}$.

- First, we assume the Stokes-Einstein relation for the rotational diffusion coefficient of agents [43]

$$D_{\mathrm{rot}}\,[\mathrm{s}^{-1}] = \frac{k_B T}{8\pi\eta\, a\,[\mu\mathrm{m}]^3} \quad , \tag{S15}$$

  where $\eta$ is the dynamic viscosity of water. This corresponds to the conservative estimate that thermal fluctuations dominate the effective rotational diffusion coefficient of motile cells, which provides a lower bound. The rescaling of time then gives $D_{\mathrm{rot}} = D_{\mathrm{rot}}\,[\mathrm{s}^{-1}]\,T$.

- Second, we employ the Berg-Purcell result for the binding rate of diffusing molecules of concentration $c$ to a spherical absorber of radius $a$ [15]

$$r_0\,[\mathrm{s}^{-1}] = \lambda\,[\mathrm{nM}^{-1}\,\mathrm{s}^{-1}]\,c_0\,[\mathrm{nM}] = 4\pi\, D_c\,[\mu\mathrm{m}^2/\mathrm{s}]\,a\,[\mu\mathrm{m}]\,c_0\,[\mathrm{nM}] \quad . \tag{S16}$$

  Here, $D_c$ denotes the translational diffusion coefficient of the signaling molecules, which we compute according to the Stokes-Einstein equation as

$$D_c\,[\mu\mathrm{m}^2/\mathrm{s}] = \frac{k_B T}{6\pi\eta\, R_c} \quad , \tag{S17}$$

  where $R_c$ is the (effective) radius of the diffusing signaling molecules. The rescaling of time and concentration then gives $\lambda c_0 = J_0\,[\mathrm{s}^{-1}]\,T$. As reference concentration, we use the concentration $c_0 = c(R_0) = 1/R_0$ at the start position, hence $c_0\,[\mathrm{nM}] = c_0 C$.

We assume the following parameters, $T = 300\,\mathrm{K}$, $\eta = 8.94 \cdot 10^{-4}\,\mathrm{kg\,m^{-1}\,s^{-1}}$, $R_c = 0.3\,\mathrm{nm}$, which gives $D_c \approx 725\,\mu\mathrm{m}^2\,\mathrm{s}^{-1}$. Furthermore, we use as typical concentration scale $c_0\,[\mathrm{nM}] = 1\,\mathrm{nM}$.

Using these assumption, we can re-plot the data from [2] as function of our empirical power-law for the relative importance of spatial comparison as shown in Fig. 4B.

### D. ADDITIONAL SIMULATIONS

#### 1. Probability of the agent to eventually find the target

The probability to find the target within time $t$ saturates to a maximal value $P_{\mathrm{reached}}$ equal to the probability to eventually find the target due to our use of absorbing boundary conditions at a maximal target distance $R_{\mathrm{max}}$, see Fig. S2a, b. This probability $P_{\mathrm{reached}}$ collapses onto a master-curve with empirical power-law, see Fig. S2c, though success probabilities are consistently higher for the maximum-likelihood strategy.

The power-law exponents obtained for $I_\varphi^*$ and $P_{\mathrm{reached}}$, respectively, have the same signs, though the impact of motility noise is reduced for $P_{\mathrm{reached}}$. Intriguingly, the ratio of the exponents for $\lambda$ and $a$ are essentially equal (Fig. 3f: 2.8, Fig. S2c: 2.75), suggesting an equal relative importance of temporal and spatial comparison.

#### 2. Size of search domain and target size

We confirmed that similar empirical power-laws are obtained if search domain size $R_{\mathrm{max}}$ or target size $R_{\mathrm{target}}$ are changed, see Figs. S4 and S5, respectively. Note that the choice of search domain radius $R_{\mathrm{max}}$ amounts to defining a transition between an outer and inner search problem [60].

#### 3. Exponential concentration field

We confirmed that similar empirical power-laws are obtained if instead of the concentration field $c(\mathbf{r}) = 1/|\mathbf{r}|$ used in Figs. 2-5 of the main text, an exponential concentration profile was used

$$c(\mathbf{x}) = A\,\exp\left(-\frac{|\mathbf{x}| - R_{\mathrm{target}}}{\rho}\right) \quad . \tag{S18}$$

Here, we have chosen amplitude $A$ and decay length $\rho$ such that $c(R_{\mathrm{target}})$ and $c(R_{\mathrm{max}})$ take the same values at the boundary of the simulation domain for Eq. (S18) and for the usual concentration field $c(\mathbf{r}) = 1/|\mathbf{r}|$, i.e., $A = 1/R_{\mathrm{target}}$ and $\rho = (R_{\mathrm{max}} - R_{\mathrm{target}})/\ln(R_{\mathrm{max}}/R_{\mathrm{target}})$. An exponential concentration field (approximately) corresponds to the concentration field established by diffusion from a single source if signaling molecule become degraded at a constant rate. The fitted exponents for the empirical power-law obtained in additional simulations using this exponential concentration profile are reported in Table S2, with simulation results for the relative importance of SC as function of this power-law shown in Fig. S6.

#### 4. Infotaxis with inaccurate modeling of the environment

We tested the robustness of target search with the infotaxis decision rule with respect to deviations between the simulated "real" environment, and the internal model the agent has of this environment. For this, we denote parameter values assumed by the agent and used in its prediction and udpate steps by a hat, to distinguish these from the true parameter values. For example, the agent assumes a value $\widehat{D}_{\mathrm{rot}}$ for its rotational diffusion coefficient, and predicts the likelihood map $p(\mathbf{x}, t)$ according to Eq. (5) with $D_{\mathrm{rot}}$ replaced by $\widehat{D}_{\mathrm{rot}}$, while the true relative position of the target (unknown to the agent) is rotated in the egocentric map in each time-step $\Delta t$ by a random angle $\Delta\phi$ drawn from a normal distribution $\mathcal{N}(0, 2D_{\mathrm{rot}}\,\Delta t)$ whose variance is set by the true rotational diffusion coefficient $D_{\mathrm{rot}}$.

Similarly, if the agent assumes a rate constant $\widehat{\lambda}$ of molecule detection, this parameter is used in updating $p(\mathbf{x}, t)$ according to Eq. (5), while the vector-valued chemotactic signal $\mathbf{s}(t)$ is simulated using the true value $\lambda$. On a technical note, infotaxis agents over-estimate target distances for $\widehat{\lambda} > \lambda$. This prompts the use of larger distance cut-offs for likelihood maps, to avoid that likely target positions are off the map. We therefore used a cut-off distance $R_{\mathrm{max}}\,\widehat{\lambda}/\lambda$ for the simulations; for numerical speed, we also scale the space discretization $\Delta x = \Delta y$ by the same factor. Note that the radius $R_{\mathrm{max}}$ of the search domain with absorbing boundary conditions is not changed, thus comparability of results is ensured.

Fig. S7 shows that infotaxis has the best performance if their assumed $\widehat{\lambda}$ is slightly smaller than the real $\lambda$, while maximum-likelihood agents perform best if their assumed $\widehat{\lambda}$ matches the real $\lambda$, irrespective of the level of motility noise.

Fig. S9 shows that assuming a smaller rotational diffusion coefficient $\widehat{D}_{\mathrm{rot}}$ can actually improve the probability $P_{\mathrm{reached}}$ to find the target, but has almost no effect on the first passage time. In contrast, agents that assume larger values $\widehat{D}_{\mathrm{rot}}$ of $D_{\mathrm{rot}}$ (which may be considered as agents with leaky memory), require more time to find the target and, in case of infotaxis, have a lower probably to find the target, irrespective of the level of sensing noise.

Fig. S9 shows that agents can still find a source at the center of a non-radial concentration field, even if agents assume a radially symmetric concentration field. For these simulations, we used a concentration field compressed along the $y$-axis by a factor $\gamma^{1/2}$, given by

$$c(x, y) = 1/\sqrt{x^2 + \gamma\, y^2} \quad . \tag{S19}$$

The behavior of agents is qualitatively similar, with moderately reduced performance for $\gamma = 2$, and further reduced performance reflected by increased conditional first passage times and decreased success probability $P_{\mathrm{reached}}$ for compression factors of up to $\gamma = 25$. For this case of non-radially symmetric concentration fields, search performance depends on the direction of the initial position of the agent relative to the axis of compression. The reduction of search performance with increasing $\gamma$ is more pronounced for an initial position $\mathbf{x}(t = 0) = (0, -R_0)^T$ on the $y$-axis, compared to an initial position $\mathbf{x}(t = 0) = (-R_0, 0)^T$ on the $x$-axis, most likely because the steepness of concentration gradients is higher along the $y$-axis as compared to the $x$-axis.

We additionally explored a scenario of a concentration field distorted by drift. Specifically, we consider the concentration field established by diffusion from a single source with diffusion coefficient $D_0$ in the presence of drift with drift speed $v_{\mathrm{drift}}$

$$c(x, y) = \frac{1}{|\mathbf{x}|}\,\exp\left[\frac{v_{\mathrm{drift}}}{2\,D_0}\,(x - |\mathbf{x}|)\right] \quad . \tag{S20}$$

Additionally, the agent is convected by this external drift, but unlike [25] is unaware of this drift, and thus does not account for it in its prediction step. Fig. S10 shows search performance for different values of $v_{\mathrm{drift}}$ and different initial positions. We find moderately reduced performance for low drift speeds ($v_{\mathrm{drift}} = 5\%\,v$), and more pronounced reductions of search performance for drift speeds up to $v_{\mathrm{drift}} = 50\%\,v$.

### E. EXPERIMENTAL METHODS

*Robot design.* We repurposed a toy robot with omnidirectional Mecanum wheels (Keyestudio) by adding a circular array of 4 equidistant light sensors (AZDelivery module based on TEMT6000 diode from Vishay) mounted on exchangable 3D-printed frames of variable sizes. The light sensors feed into a Rasperry PI zero 2W micro-computer (Raspberry Pi Foundation) via an A/D-converter (DiyArduino module, based on ADS1256 integrated circuit from Texas Instruments Inc.), all mounted on the robot. To improve the sensitivity of the motor control, we used an PWM module (Adafruit, based on PCA9685 integrated circuit from NXP Semiconductors). The measurements from the light sensors are sampled at 10 Hz and transmitted from the Raspberry Pi to a control PC, which runs the same Bayesian inference algorithm as used for Fig. 2. Every six time-steps, the PC sends an instruction with the next move direction to the robot. (In principle, the algorithm could run on the robot's microprocessor, which would require slightly more programming effort and possibly a faster microprocessor.)

*Stimulus patterns.* Stochastic gray-scale pixel patterns were projected on the floor using a beamer via a diagonally oriented mirror, see Fig. S11. The intensity of each pixel in these patterns was drawn independently from a Poisson distribution with mean $\lambda c(x,y)\Delta t/4$, with rate constant of molecule detection $\lambda = 1$ and time-step $\Delta t = 0.01$, while $c(x,y) = 1/(x^2+y^2)^{1/2}$ denotes the radial gradient also used in simulations. The pixel with dimensionless coordinates $(x,y)$ has real coordinates $(xL, yL)$ in the projection on the floor, where we use a constant length-scale $L = 217\,\mathrm{cm}$ and constant time-scale $T = 10\,\mathrm{s}$ to convert dimensionless quantities to physical units. This projected pixel pattern was refreshed at 10 Hz, i.e., every $\Delta t T = 0.1\,\mathrm{s}$, matching the sampling rate of the robot. This ensures that subsequent sensor measurements are stochastically independent. The size of pixels was chosen as approximately $5\cdot 10^{-3}L \approx 1\,\mathrm{cm}$ such that each pixel only activates a single sensor.

*Tracking.* To determine the robot's position and orientation in the laboratory frame, a camera (EOS M100 with an EF-M 22/f2.0 objective from Cannon) connected to the control PC tracks a red and green LEDs on the robot, which serve as tracking aids.

*Search domain.* The radius $LR_{\mathrm{target}}$ of the (virtual) target and the initial target distance $LR_0$ match exactly those in table S1 for simulations in Fig. 2 after re-scaling with the length-scale $L = 217\,\mathrm{cm}$. The rectangular projection area of the beamer approximately has dimensions $0.54\,L\times L$. Correspondingly, we limit the search domain to a rectangular region $[-x_{\max}, x_{\max}]\times[-y_{\max}, y_{\max}]$ with $x_{\max} = 0.5$ and $y_{\max} = 0.27$. We impose absorbing boundary conditions at the boundary of this rectangular domain, i.e., experimental runs are terminated when the center of the robot, defined as the center of the sensor array, reaches this boundary, or, alternatively, the boundary of the virtual target at the origin of the pattern.

The egocentric likelihood map of the robot is modeled as disk-like with radius 1, with a space discretization of $\Delta x = \Delta y = 0.01$ used for its numerical update.

*Sensor array size.* We use 3 different sized mounting frames for the light sensor array with respective radii $aL =$ 1.25 cm, 3.05 cm and 4.50 cm, corresponding to an effective sensing length-scale $a$ of 0.00575, 0.014 and 0.02 length units, respectively. The agents have a speed between 2 and 3 mm/s tranlating into a speed between 0.01 and 0.015 length units per time unit.

*Estimate of rotational diffusion coefficient.* To estimate the effective rotational diffusion coefficient of the robot we designed a stereotypic motility task, during which the robot moves back and forth in a random direction for a distance of 0.04, which is repeated 20 times. The position $\mathbf{x}_j$ of the robot is recorded after each such back-and-forth manevour, counted by $j = 0,\ldots,20$. We computed the mean-square displacement (MSD) as function of $j$ as $\mathrm{MSD}(j) = \langle|\mathbf{x}_j - \mathbf{x}_0|^2\rangle$, averaging over $n = 6$ realizations. Actuator noise of the robot causes effective rotational diffusion, as well as a systematic drift (due to a weak dependence of the speed of the robot on its direction of motion). As a consequence, the MSD can be approximately described by the superposition of a term linear in $j$ due to rotational diffusion and a term quadratic in $j$ due to drift, $\mathrm{MSD}(j) \approx c\,j + d\,j^2$. We determined estimates and confidence intervals for the coefficients $c$ and $d$ by a least-square fit using the python package *iminuit*, see Fig. S12.

To relate the fitted slope $c$ of the linear component to an effective rotational diffusion coefficient $D_{\mathrm{rot}}$, we ran simulations of an Active Brownian Particle (ABP) that performs the same motility task. Specifically, the ABP follows the equation of motion

$$\dot{x} = v\,\cos\psi\ . \tag{S21}$$

$$\dot{y} = v\,\sin\psi\ , \tag{S22}$$

with direction angle $\psi(t)$ obeying

$$\dot{\psi} = \xi(t) \quad , \tag{S23}$$

where $v = 0.01$, and $\xi(t)$ denotes Gaussian white noise with $\langle \xi(t) \rangle = 0$ and $\langle \xi(t)\xi(t') \rangle = 2D_{\mathrm{rot}}\, \delta(t-t')$, where the direction angle $\psi(jT)$ at the beginning of each back-and-forth maneuver of duration $T = 0.08/v$ was chosen randomly from a uniform distribution in $[0, 2\pi]$ for $j = 0, \ldots, 20$, and $\psi$ was increased by $\pi$ at each U-turn at time $t = jT + T/2$. By computing the MSD from these simulated ABP, we obtained a look-up table that monotonically maps the rotational diffusion coefficient $D_{\mathrm{rot}}$ to the slope $c$ of the MSD. Applying this look-up table, we thus obtain an estimate for the effective rotational diffusion coefficient of the robot, as well as an error interval by error propagation of the one-standard-deviation confidence interval from the least-square fit.

Fitting the experimental data (see Fig. S12), we estimate $c = (3.75 \pm 0.07) \cdot 10^{-5}$ and $d = (1.008 \pm 0.005) \cdot 10^{-5}$. Our look-up table yield $D_{\mathrm{rot}} = 0.004\,[0 - 0.013]$.

### F. DETAILS ON ANALYTICAL THEORY

In the following, we provide additional information on the minimal model of an ideal chemotactic agent in the main text, as well as on the derivation of the analytical theory presented in Eqs. (5)-(7) in the main text, of which a first version was presented in [34].

#### 1. Notes on analytical derivation of Eq. (5), Eq. (6), Eq. (7)

Eqs. (5)-(7) in the main text are directly based on results derived in [34], though rewritten in more compact form that highlights information gain from either temporal comparison (TC) and spatial comparison (SC). As an important difference, we linearized the nonlinear equations from [34] by a simple moment-closure technique, which is a prerequisite for efficient numerical simulation. While our model comprises rotational diffusion, the original work considered translational diffusion, which prompts a straight-forward modification.

For the convenience of the reader, we include the derivation of Eqs. (5)-(7) below, following closely the derivation in [34], though with simplified notation.

*Moment-closure.* Direct application of Bayes' formula results in a nonlinear jump process for the time-evolution of the likelihood distribution $p(\mathbf{x}, t)$ that involves terms quadratic in the direction vector $\mathbf{e}$ of the vector-values measurement $\mathbf{s}(t)$, see [34]. We can linearize these terms in a way that is formally similar to the derivation of noise-induced drift terms in Itō and Stratonovich calculus. Specifically, we introduce a partial expectation value $\mathbb{E}_\theta$ that averages over the direction angle $\theta$ of the last detection event $d\mathbf{s}\, dt = \mathbf{e}_\theta$ (but not over the probability that a detection event occurred at all in the last time-step). The distribution $p(\theta)$ of this direction angle $\theta$ deviates from a uniform distribution $p(\theta) \approx (2\pi)^{-1}$ only by higher-order terms of order $a|\nabla c|$ due to the presence of the concentration gradient; these deviations can be neglected in the averaging as they contribute only higher-order terms.

For later use, we note the useful identity for arbitrary vectors $\mathbf{A}$ and $\mathbf{B}$

$$\mathbb{E}_\theta \left(\mathbf{e}_\theta \cdot \mathbf{A}\right)\left(\mathbf{e}_\theta \cdot \mathbf{B}\right) = \oint d\theta\, p(\theta)\, \left(\mathbf{e}_\theta \cdot \mathbf{A}\right)\left(\mathbf{e}_\theta \cdot \mathbf{B}\right) = \mathbf{A} \cdot \mathbf{B}/2 + \mathcal{O}(a|\nabla c|) \quad , \tag{S24}$$

which directly follows from $p(\theta) = (2\pi)^{-1} + \mathcal{O}(a|\nabla c|)$, $\oint d\theta\, \cos^2\theta = \pi$, and $\oint d\theta\, \cos\theta \sin\theta = 0$.

*Time-evolution of likelihood distribution, Eq. (5).* In a time-discrete formulation, the time evolution of the likelihood distribution $p(\mathbf{x}, t)$ is described by an alternation of a prediction and an update step in each time-step of duration $dt$. Eq. (5) is then obtained by performing the continuum limit $dt \to 0$.

The prediction step forecasts the change of $p(\mathbf{x}, t)$ due diffusion and motility-induced drift; hence, we directly obtain for the likelihood distribution $p'$ after the prediction but before the update step

$$\text{Prediction step:} \qquad p'(\mathbf{x}, t+dt) = p(\mathbf{x}, t) + dt\left[D_{\mathrm{rot}} \frac{d^2}{d\varphi^2} p(\mathbf{x}, t) + \mathbf{v} \cdot \nabla p(\mathbf{x}, dt)\right] \quad . \tag{S25}$$

For the update step, we distinguish two cases: either a detection event occurred in the last time-step, or not. If an event with $\mathbf{s}\, dt = \mathbf{e}$ occurred, Bayes' rule gives

$$\text{Update step if event:} \qquad p(\mathbf{x}, t+dt) = \frac{p(\mathbf{s}\, dt = \mathbf{e}\,|\,\mathbf{x})}{\langle p(\mathbf{s}\, dt = \mathbf{e}\,|\,\mathbf{x}) \rangle} p'(\mathbf{x}, t+dt) \quad , \tag{S26}$$

where the predicted likelihood $p'(\mathbf{x}, t+dt)$ serves as Bayesian prior, $p(\mathbf{s}\, dt = \mathbf{e}\,|\,\mathbf{x})$ is the measurement probability that the agent detects a molecule at position $\mathbf{x}_0 + a\mathbf{e}$ on its circumference if the target is positioned at $\mathbf{x}^* = \mathbf{x}_0 + \mathbf{x}$, based on the current likelihood. We have

$$p(\mathbf{s}\,dt = \mathbf{e}\,|\,\mathbf{x}) = J\,(2\pi)^{-1}\left[1 + \mathbf{e}\cdot\frac{a\nabla c}{c}\right]\,dt \tag{S27}$$

$$= \;(2\pi)^{-1}\left[J + \mathbf{e}\cdot\lambda a\nabla c\right]\,dt + \mathcal{O}(a^3) \quad , \tag{S28}$$

where we replaced $J$ by $\lambda c$ in the second term affecting only terms of order $\mathcal{O}(a^3)$. Inserting this expression for the measurement probability into (S26) and expanding to second order in $a$ yields

$$p(\mathbf{x}, t+dt) = \frac{J + \mathbf{e}\cdot\lambda a\nabla c}{\langle J\rangle + \langle \mathbf{e}\cdot\lambda a\nabla c\rangle}\,p'(\mathbf{x}, t+dt) + \mathcal{O}(a^3) \quad , \tag{S29}$$

$$= \frac{J}{\langle J\rangle}\left[1 + \mathbf{e}\cdot\frac{\lambda a\nabla c}{J} - \frac{\langle \mathbf{e}\cdot\lambda a\nabla c\rangle}{\langle J\rangle} + \underbrace{\frac{\langle \mathbf{e}\cdot\lambda a\nabla c\rangle^2}{\langle J\rangle^2}}_{\to\,\mathbb{E}_\theta\ldots} - \underbrace{\frac{\mathbf{e}\cdot\lambda a\nabla c}{\langle J\rangle}\frac{\langle \mathbf{e}\cdot\lambda a\nabla c\rangle}{\langle J\rangle}}_{\to\,\mathbb{E}_\theta\ldots} + \ldots\right]p'(\mathbf{x}, t+dt) + \mathcal{O}(a^3) \tag{S30}$$

$$\overset{(*)}{=} \frac{J}{\langle J\rangle}\left[1 + \left(\mathbf{e} - \frac{\langle a\nabla c\rangle/2}{\langle c\rangle}\right)\cdot\left(\frac{a\nabla c}{c} - \frac{\langle a\nabla c\rangle}{\langle c\rangle}\right)\right]p'(\mathbf{x}, t+dt) + \mathcal{O}(a^3) \quad . \tag{S31}$$

In step $(*)$, we replaced $J$ by $\lambda c$ in terms linear in $a$, which only affects terms of order $\mathcal{O}(a^3)$. The most crucial step, however, is to linearize the underbraced nonlinear terms in (S30) using the moment-closure equation (S24). From (S31), we readily identify the measurement terms in Eq. (5).

If the agent detected no event in the last time-step, its Bayesian update step reads

$$\text{Update step if no event:}\quad p(\mathbf{x}, t+dt) = \frac{1 - J dt}{1 - \langle J\rangle\,dt}p'(\mathbf{x}, t+dt) \tag{S32}$$

$$= \big(1 - J\,dt + \langle J\rangle\,dt\big)\,p'(\mathbf{x}, t+dt) + \mathcal{O}(dt^2) \quad , \tag{S33}$$

which gives rise to a deterministic drift term $\mathbb{E}s[(J/\langle J\rangle) - 1]$ in Eq. (5).

As a comment, Eq. (5) automatically conserves total probability mass. Moreover, the scalar measurement $s(t)$ and the vectorial measurement $\mathbf{s}(t)$ only enter as innovation terms $s - \mathbb{E}s$ and $\mathbf{s} - \mathbb{E}(\mathbf{s}\,|\,s{>}0)$ in Eq. (5), respectively, i.e. their expectation value or conditional expectation value vanish, respectively.

*First time derivative of the negative Shannon entropy, Eq. (6).* We provide a derivation of Eq. (6) in the main text (which corresponds to Eq. (2) in [34]). The expected change $\mathbb{E}\,dI$ in the negative Shannon entropy in the next infinitesimal time-interval $dt$ can be written as a sum

$$\mathbb{E}\,dI = \left(\frac{dI}{dt}\right)_{|s=0} dt + \mathbb{E}\,dI[\mathbf{s}] + \mathcal{O}(dt^2) \quad , \tag{S34}$$

where $(dI/dt)_{|s=0}$ denotes the rate of information change in the absence of detection events, and $dI[\mathbf{s}]$ denotes any additional change in information if an event occurs. Here, the expectation value $\mathbb{E}$ averages over the expected measurement $\mathbf{s}$ in the next time-interval, based on the current likelihood map $p(\mathbf{x}, t)$.

The rate of information change in the absence of detection events follows from the chain rule

$$\left(\frac{dI}{dt}\right)_{|s=0} = \int d\mathbf{x}\,\frac{\delta I}{\delta p}\,\dot{p}_{|s=0} \quad , \tag{S35}$$

where $\delta I/\delta p = \ln p + 1$. We evaluate this (S35) by treating the terms constituting $\dot{p}$ in Eq. (5) individually. When doing so, all terms contributing to $\int d\mathbf{x}\,\dot{p}$ will simplify to zero, as total probability mass is conserved, i.e. $\int d\mathbf{x}\,\dot{p} = d/dt\int \mathbf{x}\,p = 0$.

- The rotational diffusion term $D_\mathrm{rot}\partial^2 p/\partial\varphi^2$ gives rise to a term

$$\int d\mathbf{x}\;(\ln p + 1)\;D_\mathrm{rot}\frac{\partial^2 p}{\partial\varphi^2} = D_\mathrm{rot}\int d\mathbf{x}\,\frac{\partial^2 \ln p + 1}{\partial\varphi^2}\,p = D_\mathrm{rot}\left\langle\frac{\partial^2 \ln p}{\partial\varphi^2}\right\rangle \tag{S36}$$

  by double partial integration.

- The advection term $\mathbf{v}\cdot\nabla p$ gives rise to a term that simplifies to zero, where we used partial integration again

$$\int d\mathbf{x}\ (\ln p+1)\ \mathbf{v}\cdot\nabla p=-\mathbf{v}\cdot\int d\mathbf{x}\,\nabla\left[\ln p+1\right]p=-\mathbf{v}\cdot\int d\mathbf{x}\,\nabla p=0\quad . \tag{S37}$$

  This is expected, as a simple translation of a likelihood map should not change its negative Shannon entropy.

- The third term (TC term) in Eq. (5) gives rise to

$$\int d\mathbf{x}\ (\ln p+1)\ p\,\mathbb{E}s\left(\frac{J}{\langle J\rangle}-1\right)=\langle\ln p\left(\langle J\rangle-J\right)\rangle\quad , \tag{S38}$$

  where we used $\mathbb{E}s=\langle J\rangle$. This term will actually cancel with another term later.

- The fourth term (SC term) in Eq. (5) is zero if not detection event occurred; hence, the corresponding term in (S35) is zero.

We now account for the additional change in negative Shannon entropy $dI[\mathbf{s}]$ if a detection event occurred. A molecular detection event with $\mathbf{s}\,dt=\mathbf{e}$ changes $p$ by a multiplicative factor $\Lambda$ to $p\Lambda$ with

$$\Lambda=\frac{J}{\langle J\rangle}\Bigg[1+\underbrace{(\mathbf{e}-\mathbb{E}(\mathbf{s}\,|\,s{>}0))\cdot\left(\frac{a\nabla c}{c}-\frac{\langle a\nabla c\rangle}{\langle c\rangle}\right)}_{\Gamma}\Bigg]\quad , \tag{S39}$$

where we introduced the short-hand $\Gamma$ for the contribution to $\Lambda$ due to spatial comparison (SC). If no detection event occurred, we have $\mathbb{E}(dI[\mathbf{s}]\,|\,s=0)=0$ by definition. Thus,

$$\mathbb{E}\,dI[\mathbf{s}]=\underbrace{p_{\text{event}}}_{\langle J\rangle dt}\,\mathbb{E}\big(dI[\mathbf{s}]\,\big|\,s{>}0\big)\quad , \tag{S40}$$

where $p_{\text{event}}=\langle J\rangle\,dt$ is the probability that a detection event will occur in the next time-interval $dt$.

For later use, we note the conditional expectation values

$$\mathbb{E}(\Gamma\,|\,s>0)=0\quad ,\quad \text{and} \tag{S41}$$

$$\mathbb{E}(\Lambda\,|\,s>0)=J/\langle J\rangle\quad . \tag{S42}$$

We also note the conditional second moment of $\Gamma^2$

$$\mathbb{E}\left(\Gamma^2\,|\,s{>}0\right)=\int d\mathbf{e}\,p(\mathbf{s}\,dt=\mathbf{e}\,|\,s{>}0)\left|(\mathbf{e}-\mathbb{E}(\mathbf{s}\,|\,s{>}0))\cdot\left(\frac{a\nabla c}{c}-\frac{\langle a\nabla c\rangle}{\langle c\rangle}\right)\right|^2 \tag{S43}$$

$$=\oint d\theta\,p(\theta)\left|\mathbf{e}_\theta\cdot\left(\frac{a\nabla c}{c}-\frac{\langle a\nabla c\rangle}{\langle c\rangle}\right)\right|^2+\mathcal{O}(a^4) \tag{S44}$$

$$=\frac{1}{2}\left|\frac{a\nabla c}{c}-\frac{\langle a\nabla c\rangle}{\langle c\rangle}\right|^2+\mathcal{O}(a^3)\quad , \tag{S45}$$

where we used in step (S44) that the distribution $p(\theta)$ of direction angles $\theta$ of the vector-valued measurement $\mathbf{s}\,dt=\mathbf{e}_\theta$ is uniform to leading order in $a$, $p(\theta)=(2\pi)^{-1}+\mathcal{O}(a)$, as well as that $\mathbb{E}(\mathbf{s}\,|\,s{>}0)$ only contributes higher-order terms of order $\mathcal{O}(a^4)$, which reduces the calculation to the square of a direction cosine, hence the factor $1/2$.

If a detection event occurs, and $p$ changes to $p\Lambda$, the negative Shannon entropy of $p$ changes by an amount $dI[\mathbf{s}]$

$$\begin{aligned} dI[\mathbf{s}] &= I\left[p\Lambda\right]-I[p] \\ &= \int d\mathbf{x}\,p\Lambda\ln\left(p\Lambda\right)-\int d\mathbf{x}\,p\ln p \\ &= \int d\mathbf{x}\,p\left(\Lambda-1\right)\ln p+\int d\mathbf{x}\,p\Lambda\ln\Lambda\quad . \end{aligned} \tag{S46}$$

Taking the conditional expectation value of the first term in (S46) yields [because of (S42)]

$$\mathbb{E}\bigg(\int d\mathbf{x}\,p\left(\Lambda-1\right)\ln p\,\bigg|\,s>0\bigg)=\left\langle\left(\frac{J}{\langle J\rangle}-1\right)\ln p\right\rangle\quad . \tag{S47}$$

This term multiplied by $p_{\text{event}} = \langle J\rangle\, dt$ according to (S40) cancels with the contribution from (S38) in (S34).

We now compute the contribution of the second term in (S46) to (S40), by taking its conditional expectation value and multiplying with $p_{\text{event}} = \langle J\rangle\, dt$

$$\langle J\rangle\, dt\, \mathbb{E}\Big(\int d\mathbf{x}\, p\Lambda \ln\Lambda \,\Big|\, s>0\Big) \tag{S48}$$

$$= \langle J\rangle\, dt\, \mathbb{E}\Big\{\int d\mathbf{x}\, p\frac{J}{\langle J\rangle}(1+\Gamma)\left(\ln\left(\frac{J}{\langle J\rangle}\right)+\ln(1+\Gamma)\right)\,\Big|\, s>0\Big\} \tag{S49}$$

$$= dt\int d\mathbf{x}\, p\, J\, \mathbb{E}(1+\Gamma\,|\, s>0)\ln\left(\frac{J}{\langle J\rangle}\right) + dt\int d\mathbf{x}\, p\, J\, \mathbb{E}\big[(1+\Gamma)\left(\Gamma-\Gamma^2/2+\ldots\right)\,|\, s>0\big] \tag{S50}$$

$$= \left\langle J\ln\left(\frac{J}{\langle J\rangle}\right)\right\rangle dt + \big\langle J\, \mathbb{E}(\Gamma^2\,|\, s>0)/2\big\rangle\, dt + \mathcal{O}(a^3) \quad . \tag{S51}$$

These two terms constitute the second term (TC term) and third term (SC term) of Eq. (6), respectively [using (S45)].

*Second time derivative of the negative Shannon entropy, Eq. (7).* Auconi et al. gives a result for the second variation of the entropy $S$ in their Supplemental Materials, section IV.B, see Eq. (36) there [34]. However, this previous result is convoluted and its physical significance maybe difficult to extract. We therefore re-derive Eq. (7) from Eq. (6) directly, building on the previous simplifications.

We start with the 'chain rule' for the case that no detection event occurred in the infinitesimal time-interval $dt$

$$\frac{d}{dt}\left(\mathbb{E}\,\frac{dI}{dt}\right)_{|s=0} = \int d\mathbf{x}\,\left(\frac{\delta}{\delta p}\,\mathbb{E}\,\frac{dI}{dt}\right)\dot{p}_{s=0} = \int d\mathbf{x}\,\left(\frac{\delta}{\delta p}\,\mathbb{E}\,\frac{dI}{dt}\right)\nabla p\cdot\mathbf{v}+\ldots \quad , \tag{S52}$$

where we used Eq. (5) and omitted terms independent of $\mathbf{v}$. If we account for the additional change in $p$ in the case of a detection event, we obtain additional terms, but all these terms are independent of $\mathbf{v}$ and hence can be omitted. Thus, independent of the value of $s\, dt$ in this time-interval, we can write, using partial integration

$$\frac{d}{dt}\left(\mathbb{E}\,\frac{dI}{dt}\right) = -\mathbf{v}\cdot\nabla\left(\frac{\delta}{\delta p}\,\mathbb{E}\,\frac{dI}{dt}\right)+\ldots \quad , \tag{S53}$$

where terms independent of $\mathbf{v}$ have been omitted.

We thus need to compute functional derivatives. For the first term of Eq. (6), we find

$$\frac{\delta}{\delta p}\,\langle J\ln J\rangle = J\ln J \tag{S54}$$

and

$$\frac{\delta}{\delta p}\,\langle J\ln\langle J\rangle\rangle = J\ln\langle J\rangle + J \quad . \tag{S55}$$

Here, the additional term $J = \langle J\rangle/\langle J\rangle J$ arises from taking the variation of the inner average $\langle J\rangle$ in $J\ln\langle J\rangle$. Combining Eqs. (S54) and (S55) gives

$$\nabla\frac{\delta}{\delta p}\left[J\ln\left(\frac{J}{\langle J\rangle}\right)\right] = \nabla[J\ln J - J\ln\langle J\rangle - J] = (\nabla J)\ln\left(\frac{J}{\langle J\rangle}\right) \quad . \tag{S56}$$

For the second term of Eq. (6) in the main text, we first note

$$\left\langle c\left|\frac{\nabla c}{c}-\frac{\langle\nabla c\rangle}{\langle c\rangle}\right|^2\right\rangle = \left\langle\frac{|\nabla c|^2}{c}\right\rangle - 2\frac{|\langle\nabla c\rangle|^2}{\langle c\rangle} + \langle c\rangle\,\frac{|\langle\nabla c\rangle|^2}{\langle c\rangle^2} = \underbrace{\left\langle\frac{|\nabla c|^2}{c}\right\rangle}_{=a} - \underbrace{\frac{|\langle\nabla c\rangle|^2}{\langle c\rangle}}_{=b} \quad , \tag{S57}$$

from which we can compute the functional derivative (using the chain rule)

$$\frac{\delta}{\delta p}\left\langle c\left|\frac{\nabla c}{c}-\frac{\langle\nabla c\rangle}{\langle c\rangle}\right|^2\right\rangle = \underbrace{\frac{|\nabla c|^2}{c}}_{\partial a/\partial p} \underbrace{-2\frac{\langle\nabla c\rangle\cdot\nabla c}{\langle c\rangle}+\frac{|\langle\nabla c\rangle|^2}{\langle c\rangle^2}c}_{\partial b/\partial p} = c\left|\frac{\nabla c}{c}-\frac{\langle\nabla c\rangle}{\langle c\rangle}\right|^2 \quad . \tag{S58}$$

Multiplying this result with $a^2/4$ and inserting it into (S53) yields, up to terms of order $\mathcal{O}(a^4)$, the second term of Eq. (7) in the main text.

### 2. Relation to Auconi et al. [34]

To facilitate a comparison of the original mathematical theory in [34], and the adapted version comprising Eqs. (5)-(7) in the main text, we show how these equations map to the original analytical results in [34].

*Notation.* We start with a comparison of the notation used here and in [34]. Auconi et al. used a notation akin to Wiener processes to describe the stochastic update dynamics, introducing a cumulative measurement process $\mathbf{m}(t) = \int_0^t d\mathbf{m}(t) = \int_0^t dt\, \mathbf{s}(t)$; for the increments, we can use the short-hand notation $d\mathbf{m} = d\mathbf{s}\, dt$, where $dt$ denotes an infinitesimal time-step. Further, in this work, we write $J(\mathbf{x})$ for the rate of molecule detection by an agent of size $a$ if the target is located at relative position $\mathbf{x}$, while [34] used this symbol for the corresponding rate $\lambda c(\mathbf{x})$ of a point-like agent. In the following, we will write $J_0(\mathbf{x}) = \lambda c(\mathbf{x})$ for this rate to avoid ambiguity. All changes of notation are summarized in table S3.

| Notation used in this work | Notation used in Auconi et al. [34] | Meaning |
|---|---|---|
| $\mathbf{x}^*$ | $\mathbf{x}$ | target position |
| $\mathbf{x}_0$ | $\mathbf{0}$ | position of agent |
| $\mathbf{x}(t)$ | $\mathbf{x}_0$ | relative target position *before* update step |
| $\mathbf{e}$ | $e^{\mathrm{i}\theta}$ | unit vector |
| $\mathbf{x_e} = \mathbf{x}_0 + a\mathbf{e}$ | $ae^{\mathrm{i}\theta}$ | point on agent's circumference |
| $\lambda c(\mathbf{x})$ | $r(\mathbf{x})$ | detection rate of point-agent |
| $\lambda c(\mathbf{x_e})/(2\pi)$ | $r(\mathbf{x} - a\, e^{\mathrm{i}\theta})/(2\pi)$ | detection rate on circumference in direction $\mathbf{e}$ |
| $J(\mathbf{x}) \approx \lambda[c(\mathbf{x}) + (a^2/4)\nabla^2 c(\mathbf{x})]$ | $r(\mathbf{x}) + (a^2/4)\nabla^2 r(\mathbf{x})$ | detection rate of agent of size $a$ |
| $\mathbf{s}\, dt$ | $d\mathbf{m}$ | increment of measurement process |
| $s\, dt$ | $dm = \lvert d\mathbf{m}\rvert^2$ | increment of scalar measurement process |
| $\nabla c = \nabla_{\mathbf{x}_0} c(\mathbf{x}_0\lvert\mathbf{x}^*)$ | $-\nabla r/\lambda$ | concentration gradient |
| $\nabla J = \nabla_{\mathbf{x}_0} J(\mathbf{x}_0\lvert\mathbf{x}^*)$ | $-\nabla[r + (a^2/4)\nabla^2 r]$ | gradient of effective detection rate |
| $\nabla p = \nabla_{\mathbf{x}} p$ | $\nabla p$ | gradient of likelihood map |
| $dt$ | $\tau$ | time-step |
| $I$ | $-S$ | negative Shannon entropy |

TABLE S3. Comparison of notations used in this work and in [34].

*Time-evolution of the likelihood distribution.* In the limit of a point-like agent, the first three summands in our Eq. (5) correspond exactly to Eq. (13) in section I.A of the Supplementary Materials of [34]. Note that $\mathbb{E}\, s = J$; hence, $s - \mathbb{E}\, s$ in our notation corresponds to $dm - (J_0 + a^2\nabla^2 J_0/4)\, dt$ in the notation of [34]. As we consider rotational diffusion in our model, the diffusion term is replaced by $D_{\mathrm{rot}}\partial^2 p/\partial\varphi^2$, where $\varphi$ denotes the azimuthal angle of polar coordinates, which directly follows from the representation of the Laplace operation in polar coordinates. Except for the diffusion, this result is similar to the analytic result derived in [27].

The derivation of terms resulting from the finite size $a$ of agents in our Eq. (5) is based on Eq. (28) in section III of the Supplementary Materials of [34]. It is this step, where we perform the linearization of the originally nonlinear Itō evolution equation for the likelihood $p$. For the convenience of the reader, we quote said Eq. (28)

$$\begin{aligned}\frac{dp}{p} = &\left[D\frac{\nabla^2 p}{p} + \nabla \ln p \cdot \mathbf{v} + \langle J_0\rangle - J_0 + \frac{a^2}{4}\left(\langle\nabla^2 J_0\rangle - \nabla^2 J_0\right)\right] dt \\ &+ \left(\frac{J_0}{\langle J_0\rangle} - 1\right) \lvert d\mathbf{m}\rvert^2 + a\,\frac{J_0}{\langle J_0\rangle}\left(\frac{\langle\nabla J_0\rangle}{\langle J_0\rangle} - \frac{\nabla J_0}{J_0}\right)\cdot d\mathbf{m} \\ &+ \frac{a^2}{2}\frac{J_0}{\langle J_0\rangle}\left[d\mathbf{m}\cdot\left(\frac{\mathbf{H}}{J_0} - \frac{\langle\mathbf{H}\rangle}{\langle J_0\rangle}\right)\cdot d\mathbf{m} + 2\left(d\mathbf{m}\cdot\frac{\langle\nabla J_0\rangle}{\langle J_0\rangle}\right)\left(d\mathbf{m}\cdot\left[\frac{\langle\nabla J_0\rangle}{\langle J_0\rangle} - \frac{\nabla J_0}{J_0}\right]\right)\right] \quad . \end{aligned} \tag{S59}$$

The steps of rewriting this equation are

- The diffusion term for translational diffusion is replaced by the corresponding term for rotational diffusion

$$D\nabla^2 p \to D_{\mathrm{rot}}\frac{\partial^2}{\partial\varphi^2}p \quad , \tag{S60}$$

which directly follows from the representation of the Laplace operation in polar coordinates with polar angle $\varphi$.

- The advection term can be readily rewritten

$$p\nabla \ln p \cdot \mathbf{v} = \mathbf{v}\cdot\nabla p \quad . \tag{S61}$$

- For the next term, we first note

$$\langle J_0\rangle - J_0 + \frac{a^2}{4}\left(\langle\nabla^2 J_0\rangle - \nabla^2 J_0\right) = \langle J\rangle - J + \mathcal{O}(a^3) \quad , \tag{S62}$$

which is equal to $-p\mathbb{E}s[(J/\langle J\rangle) - 1]$ because of $\mathbb{E}s = \langle J\rangle$. Next, using $|d\mathbf{m}|^2 = s\,dt$, we identify the term $p[(J_0/\langle J_0\rangle - 1]|d\mathbf{m}|^2$ in (S59). This term agrees to leading order with the term $ps[J/\langle J\rangle - 1]$ in our Eq. (5) since

$$\frac{J}{\langle J\rangle} = \frac{J_0}{\langle J_0\rangle} + \frac{a^2}{4}\left[\frac{\nabla^2 J_0}{\langle J_0\rangle} - J_0\frac{\langle\nabla^2 J_0\rangle}{\langle J_0\rangle^2}\right] + \mathcal{O}(a^3) \quad . \tag{S63}$$

Last, we linearize a term quadratic in the measurement increment $d\mathbf{m}$

$$d\mathbf{m}\cdot\left(\frac{\mathbf{H}}{J_0} - \frac{\langle\mathbf{H}\rangle}{\langle J_0\rangle}\right)\cdot d\mathbf{m} \tag{S64}$$

by replacing this term with its partial expectation value $\mathbb{E}_\theta(d\mathbf{m}\cdot\mathbf{H}\cdot d\mathbf{m})$. We thus obtain

$$\mathbb{E}_\theta(d\mathbf{m}\cdot\mathbf{H}\cdot d\mathbf{m}) = \oint d\theta\, p(\theta)\, d\mathbf{m}\cdot\mathbf{H}\cdot d\mathbf{m} = s\,dt\,\frac{\operatorname{tr}\mathbf{H}}{2} + \mathcal{O}(a) = s\,dt\,\frac{\nabla^2 J_0}{2} + \mathcal{O}(a) \quad . \tag{S65}$$

Here, we used that $s\,dt = dm$ can only take values 0 or 1 in a short measurement time interval, hence $dm^2 = dm$, as well as $\operatorname{tr}\mathbf{H} = \nabla^2 J_0$ for the Hessian $\mathbf{H}$ of $J_0$. Applying this approximation to (S59) yields

$$\frac{a^2}{2}\frac{J_0}{\langle J_0\rangle}d\mathbf{m}\cdot\left(\frac{\mathbf{H}}{J_0} - \frac{\langle\mathbf{H}\rangle}{\langle J_0\rangle}\right)\cdot d\mathbf{m} \to s\,dt\,\frac{a^2}{4}\left(\frac{\nabla^2 J_0}{\langle J_0\rangle} - J_0\frac{\langle\nabla^2 J_0\rangle}{\langle J_0\rangle^2}\right) \quad . \tag{S66}$$

By (S63), we have thus collected all subterms of the third term in Eq. (5), $p(s - \mathbb{E}\,s)[(J/\langle J\rangle) - 1]$.

- As $d\mathbf{m} = \mathbf{s}\,dt$ and $\nabla c = -\nabla J_0$, we readily notice

$$a\,\frac{J_0}{\langle J_0\rangle}\left(\frac{\langle\nabla J_0\rangle}{\langle J_0\rangle} - \frac{\nabla J_0}{J_0}\right)\cdot d\mathbf{m} \to \frac{J_0}{\langle J_0\rangle}\,\mathbf{s}\cdot\left(\frac{a\nabla c}{c} - \frac{\langle a\nabla c\rangle}{\langle c\rangle}\right)dt \quad . \tag{S67}$$

Replacing the factor $J_0/\langle J_0\rangle$ by $J/\langle J\rangle$ only introduces higher-order corrections $\mathcal{O}(a^3)$. Next, we linearize the last, nonlinear term of (S59), using moment-closure, using an analogous procedure as in the previous step. Applying the identity (S24) with $\mathbf{A}=\langle\nabla J_0\rangle/\langle J_0\rangle$ and $\mathbf{B}=\langle\nabla J_0\rangle/\langle J_0\rangle - \nabla J_0/J_0$, yields

$$\mathbb{E}_\theta\left\{2\left(d\mathbf{m}\cdot\frac{\langle\nabla J_0\rangle}{\langle J_0\rangle}\right)\left(d\mathbf{m}\cdot\left[\frac{\langle\nabla J_0\rangle}{\langle J_0\rangle} - \frac{\nabla J_0}{J_0}\right]\right)\right\} = s\,dt\,\frac{\langle\nabla J_0\rangle}{\langle J_0\rangle}\cdot\left(\frac{\langle\nabla J_0\rangle}{\langle J_0\rangle} - \frac{\nabla J_0}{J_0}\right) \quad . \tag{S68}$$

We thus note the replacement

$$\frac{a^2}{2}\frac{J_0}{\langle J_0\rangle}2\left(d\mathbf{m}\cdot\frac{\langle\nabla J_0\rangle}{\langle J_0\rangle}\right)\left(d\mathbf{m}\cdot\left[\frac{\langle\nabla J_0\rangle}{\langle J_0\rangle} - \frac{\nabla J_0}{J_0}\right]\right) \to -\frac{J_0}{\langle J_0\rangle}s\frac{\langle\lambda a\nabla_{\mathbf{x}_0}c/2\rangle}{\langle J_0\rangle}\cdot\left(\frac{a\nabla_{\mathbf{x}_0}c}{c} - \frac{\langle a\nabla_{\mathbf{x}_0}c\rangle}{\langle c\rangle}\right)dt \quad , \tag{S69}$$

where we used that $a\nabla_{\mathbf{x}}J_0/2 = -\lambda a\nabla_{\mathbf{x}_0}c/2 = -\mathbb{E}\mathbf{s}$. With $\mathbb{E}(\mathbf{s}\,|\,s > 0) = \mathbb{E}\mathbf{s}\,s/\langle J\rangle = \mathbb{E}\mathbf{s}\,s/\langle J_0\rangle + \mathcal{O}(a^2)$, and $J/\langle J\rangle = J_0/\langle J_0\rangle + \mathcal{O}(a^2)$, we collected all subterms of the fourth term in Eq. (5) in the main text.

The linearization step (S66) amounts to a moment-closure for the nonlinear jump process (S59).

*First time derivative of the negative Shannon entropy.* Our Eq. (6) for the temporal change of the negative Shannon entropy $I[p(\mathbf{x})] = \int d\mathbf{x}\, \ln p(\mathbf{x})$ corresponds to Eq. (2) in [34], again with the term corresponding to diffusion replaced by the analogous term for rotational diffusion in our case.

For sake of reference, we quote Eq. (2) from [34] (note that the detection rate $J_0$ of a point-like agent was denoted $r$ there)

$$\frac{\langle dS\rangle}{dt} = -D\left\langle \nabla^2 \ln p\right\rangle + \left\langle \left(r_0 + \frac{a^2}{4}\nabla^2 J_0\right)\ln\left(\frac{\langle J_0\rangle}{J_0}\right)\right\rangle + \frac{a^2}{4}\left(\frac{|\langle \nabla J_0\rangle|^2}{\langle J_0\rangle} - \left\langle \frac{|\nabla J_0|^2}{J_0}\right\rangle\right) + \mathcal{O}(a^3) \quad . \tag{S70}$$

The factor $J_0 + (a^2/4)\nabla^2 J_0$ in the second term is readily identified as our $J$. An explicit calculation shows that

$$\ln\left(\frac{J}{\langle J\rangle}\right) = \ln\left(\frac{J_0}{\langle J_0\rangle}\right) + \mathcal{O}(a^4) \tag{S71}$$

because the second-order term $\left\langle J_0(\nabla^2 J_0/J_0)\right\rangle - \left\langle \left(\left\langle \nabla^2 J_0\right\rangle / \langle J_0\rangle\right) J_0\right\rangle$ of a Taylor expansion in $a$ simplifies to zero.

For the last term in Eq. (2) of [34] (that is, (S70)), we can apply a step similar to (S57)

$$\begin{aligned}
-\frac{a^2}{4}\left(\frac{|\langle \nabla J_0\rangle|^2}{\langle J_0\rangle} - \left\langle \frac{|\nabla J_0|^2}{J_0}\right\rangle\right) &= \frac{a^2}{4}\left\langle r_0\left(-\frac{|\langle \nabla J_0\rangle|^2}{\langle J_0\rangle^2} + \frac{|\nabla J_0|^2}{J_0^2}\right)\right\rangle \\
&= \frac{a^2}{4}\left\langle J_0\left(+\frac{|\langle \nabla J_0\rangle|^2}{\langle J_0\rangle^2} - 2\frac{\langle \nabla J_0\rangle\cdot\nabla J_0}{\langle J_0\rangle J_0} + \frac{|\nabla J_0|^2}{J_0^2}\right)\right\rangle &\text{(S72)}\\
&= \frac{a^2}{4}\left\langle J_0\left|\frac{\langle \nabla J_0\rangle}{\langle J_0\rangle} - \frac{\nabla J_0}{J_0}\right|^2\right\rangle &\text{(S73)}\\
&= \frac{1}{4}\left\langle J\left|\frac{a\nabla c}{c} - \frac{\langle a\nabla c\rangle}{\langle c\rangle}\right|^2\right\rangle + \mathcal{O}(a^4) \quad . &\text{(S74)}
\end{aligned}$$

The original derivation of Eq. (2) of [34] can be found in section II.A of its Supplementary Materials, especially their Eq. (20), for the terms corresponding to a point-like agent, as well as section IV.A, especially their Eq. (35) for the additional terms due to the finite size $a$ of the agent.

*Second time derivative of the negative Shannon entropy.* We show that our Eq. (7) is equivalent to the result in [34]. For a point-like agent, our Eq. (7) reduces to Eq. (23) in section I.B in the Supplementary Materials of [34]. Additional terms due to the finite size $a$ of the agent are derived in section IV.B of [34]. We quote the relevant Eq. (36) from [34]

$$\begin{aligned}
d\langle dS\rangle &= \langle (\ln J_0 - \ln\langle J_0\rangle)\,\nabla J_0\rangle\cdot\mathbf{v}\,(dt)^2 \\
&+ \frac{a^2}{4}\Bigg[\underbrace{\left\langle \nabla\nabla^2 J_0 \ln J_0\right\rangle}_{(a)} - \underbrace{\left\langle \nabla\nabla^2 J_0\right\rangle \ln\langle J_0\rangle}_{(b)} + \underbrace{\left\langle \frac{(\nabla^2 J_0)\nabla J_0}{J_0}\right\rangle}_{(c)} - \underbrace{\frac{\left\langle \nabla^2 J_0\right\rangle\langle \nabla J_0\rangle}{\langle J_0\rangle}}_{(d)} - \underbrace{2\frac{\langle \nabla J_0\rangle}{\langle J_0\rangle}\cdot\langle \nabla\nabla J_0\rangle}_{(e)} + \underbrace{2\left\langle \frac{\nabla J_0}{J_0}\cdot\nabla\nabla J_0\right\rangle}_{(f)} \\
&\qquad + \underbrace{\frac{|\langle \nabla J_0\rangle|^2\langle \nabla J_0\rangle}{\langle J_0\rangle^2}}_{(g)} - \underbrace{\left\langle \frac{|\nabla J_0|^2\nabla J_0}{J_0^2}\right\rangle}_{(h)}\Bigg]\cdot\mathbf{v}\,(dt)^2 + \ldots,
\end{aligned} \tag{S75}$$

where terms independent of velocity $\mathbf{v}$ were omitted.

The factor of the first term (TC term) in our Eq. (7) can be expanded as

$$\left\langle \ln\left(\frac{J}{\langle J\rangle}\right)\nabla_{\mathbf{x}_0} J\right\rangle = -\left\langle \ln\left(\frac{J}{\langle J\rangle}\right)\nabla_{\mathbf{x}} J_0\right\rangle - \frac{a^2}{4}\left\langle \ln\left(\frac{J}{\langle J\rangle}\right)\nabla_{\mathbf{x}}\nabla^2 J\right\rangle \quad , \tag{S76}$$

and further

$$\left\langle \ln\left(\frac{J}{\langle J\rangle}\right)\nabla J_0\right\rangle = \langle (\ln J_0 - \ln\langle J_0\rangle)\,\nabla J_0\rangle + \frac{a^2}{4}\Bigg[\underbrace{\left\langle \frac{(\nabla^2 J_0)\nabla J_0}{J_0}\right\rangle}_{\to(c)} - \underbrace{\frac{\left\langle \nabla^2 J_0\right\rangle\langle \nabla J_0\rangle}{\langle J_0\rangle}}_{\to(d)}\Bigg] + \mathcal{O}(a^4) \quad , \tag{S77}$$

as well as

$$\frac{a^2}{4}\left\langle \ln\left(\frac{J}{\langle J\rangle}\right)\nabla\nabla^2 J\right\rangle = \frac{a^2}{4}\Bigg[\underbrace{\langle\nabla\nabla^2 J_0 \ln J_0\rangle}_{\to(a)} - \underbrace{\langle\nabla\nabla^2 J_0\rangle \ln\langle J_0\rangle}_{\to(b)}\Bigg] \quad . \tag{S78}$$

Note the sign changes $\nabla_{\mathbf{x}_0} = -\nabla_{\mathbf{x}}$ and $I = -S$.

The factor of the second term (SC term) in our Eq. (7) can be expanded as

$$\left\langle \nabla_{\mathbf{x}_0}\left(\frac{J}{4}\left|\frac{a\nabla c}{c} - \frac{\langle a\nabla c\rangle}{\langle c\rangle}\right|^2\right)\right\rangle = -\frac{a^2}{4}\left\langle \nabla_{\mathbf{x}}\left(J_0\left|\frac{\nabla J_0}{J_0} - \frac{\langle\nabla J_0\rangle}{\langle J_0\rangle}\right|^2\right)\right\rangle + \mathcal{O}(a^4) \tag{S79}$$

$$= -\frac{a^2}{4}\left\langle \nabla_{\mathbf{x}}\left(\frac{|\nabla J_0|^2}{J_0} - \underbrace{2\frac{\nabla J_0\,\langle\nabla J_0\rangle}{\langle J_0\rangle}}_{\to\ (e)} + \underbrace{\frac{J_0|\,\langle\nabla J_0|^2\rangle}{\langle J_0\rangle^2}}_{\to\ (g)}\right)\right\rangle + \mathcal{O}(a^4) \quad . \tag{S80}$$

For the first term, we note using the product rule

$$-\frac{a^2}{4}\left\langle \nabla_{\mathbf{x}}\left(\frac{|\nabla J_0|^2}{J_0}\right)\right\rangle = -\frac{a^2}{4}\left\langle \underbrace{2\frac{\nabla J_0\cdot\nabla\nabla J_0}{J_0}}_{\to\ (f)} - \underbrace{\frac{|\nabla J_0|^2}{J_0^2}\nabla J_0}_{\to\ (h)}\right\rangle \quad . \tag{S81}$$

#### 3. Relation to original infotaxis model by Vergassola et al. [25]

In the main text, we consider an ideal chemotactic agent that employs either a maximum-likelihood or an infotaxis strategy. Compared to the original infotaxis model introduced by Vergassola et al. [25], the agents considered here can move in arbitrary directions as in [27]. Additionally, they are subject to rotational diffusion with effective rotational diffusion coefficient $D_{\mathrm{rot}}$ to model motility noise [40, 61] which can be efficiently implemented using an egocentric likelihood map in which the agent is at rest, as compared to the allocentric map originally used in [25]. The implementation of rotational diffusion is the computationally most expensive step in our simulations. The decision incentive used in [25] comprises an exploration as well as an exploitation term, which there accounts for the possibility to find the target in the next time step (see their Eq. (11)). Our Eq. (7) only comprises an analogous exploration term but lacks the exploitation term from Vergassola et al. [25]. We confirmed in preliminary numerical simulations that a corresponding exploitation term is more than three orders of magnitude smaller than the exploration term, and can hence be neglected without changing results. While previous work focused on a regime of rare molecule detections (low $\lambda$), our work demonstrates the usefulness of infotaxis also for higher rates of molecule detection (intermediate to large $\lambda$), in particular when information is continuously lost due to rotational diffusion. Finally, our agents are spatially extended and hence can perform spatial comparison in addition to temporal comparison, which marks the main novelty compared to [25, 27, 31].

### G. RELATION TO PARTIALLY OBSERVABLE MARKOV DECISION PROCESSES

There exist a rich literature on partially observable Markov decision processes (POMDP) [33, 62, 63], which provides an alternative formulation for olfactory search problems [29–32, 57]. To connect our work to this existing literature, we restate our model using the language of POMDPs below.

In POMDPs, a *state* $s_t \in \mathcal{S}$ in a set of states $\mathcal{S}$ evolves according to a stochastic Markovian dynamics with transition probabilities $p(s_{t+dt}|s_t, a_{t+dt})$ that depend on *actions* $a_{t+dt} \in \mathcal{A}$ chosen by the agent at each time step from a set of possible actions $\mathcal{A}$. For the ideal chemotactic agent considered in this work, $s_t$ would be the relative position $\mathbf{x}^*$ of the target in the egocentric frame of reference of the chemotactic agent at time $t$, while $a_{t+dt} \in [0, 2\pi)$ would be the angular direction of the velocity $\mathbf{v}$ in the next movement step, see table S4.

The agent is unaware of the true state $s_t$, and receives only partial information in the form of an observation $o_t$ correlated with $s_t$ at each time step. This enables the agent to update a *belief* $b_t(s)$ of the current state as a probability distribution over $\mathcal{S}$ according to Bayes' formula

$$b_{t+dt}(s) = \frac{p(o_t|s)}{\sum_{s\in\mathcal{S}} p(o_t|s)\, b(s)} b_t(s) \quad , \tag{S82}$$

with obvious adjustments in case of an additional prediction step, analogous to our Eqs. (S25, S26, S33). In our case, observations amount to directional molecular detection events on the circumference of the chemotactic agent, with rates determined by a concentration field with the target as source, which generalize the *hits* considered in ref. [32]. The agent aims to reach special terminal states $\mathcal{S}^\Omega \subset \mathcal{S}$, which correspond to finding the target in our case. The agent employs a *policy* $a_{t+dt} = \pi(b_t) \in \mathcal{A}$ to select the next action based on the current belief $b_t$. The task of the POMDP is to find the optimal policy $\pi^*$ that maximizes the expectation value $v^* = v_{\pi^*} = \max_\pi v_\pi$ of an objective function $v_\pi$, such as minus the mean-first-passage time (MFPT) to reach a terminal state, $v_\pi(b_0)$, for initial states distributed according to the belief $b_0$.

The *Bellman equation* states a necessary condition for the optimal value function

$$v^*(b) = -dt + \max_{a\in\mathcal{A}} \sum_{b'} P(b'|b,a)\, v^*(b') \quad . \tag{S83}$$

For time-continuous problems, the Bellman optimality equation becomes the *Hamilton-Jacobi-Bellman equation* for a stochastic optimal control problem.

While polynomial-time algorithms exist to determine optimal policies for Markov decision processes, where agents have full knowledge of the current state [64], no efficient algorithms exist for POMDPs, and solving the Bellman equation can be prohibitively difficult except for the simplest cases. Instead, various heuristics including infotaxis [25, 33], or more recently, deep-reinforcement learning [32], were used to find approximate solutions. Infotaxis turned out to be surprisingly reliable, efficient, and safe [31].

Finally, despite its elegance, we remark that the MFPT might not be the ideal metric in all application cases. For search problems in unbounded space, the distribution of first-passage time can possess power-law tails, which can confound the MFPT as most suitable metric. Additionally, if the failure probability to miss the target is larger than zero as in our case, the task turns into a multi-objective optimization problem to minimize both the failure probability and a conditional MFPT, which requires the choice of a heuristic trade-off parameter.

| Terminology for POMDP | Terminology used in this work |
|---|---|
| state $s$ | true target position $\mathbf{x}^*$ |
| belief $b$ | likelihood map of target position $p(\mathbf{x},t)$ |
| hit $o$ | molecular detection event $\mathbf{e}_j\delta(t-t_j)$ |
| policy $\pi$ | decision rule (here: *infotaxis* or *maximum-likelihood*) |
| action $a_t$ | movement step with velocity $\mathbf{v}$ |

TABLE S4. Comparison of terminology commonly used for partially observable Markov decision processes (POMDP) and terminology used in this work.

| Symbol | Description |
|---|---|
| $a$ | agent size |
| $\mathbf{x}_0$ | agent position |
| $\mathbf{x}^*$ | target position |
| $\mathbf{x}$ | relative target position $\mathbf{x} = \mathbf{x}^* - \mathbf{x}_0$ |
| $\mathbf{v}$ | velocity vector of agent |
| $v$ | speed of agent |
| $D_{\mathrm{rot}}$ | rotational diffusion coefficient of agent |
| $c(\mathbf{x}_0\|\mathbf{x}^*)$ | concentration field of signaling molecules, provided target at $\mathbf{x}^*$ |
| $\lambda$ | rate constant of molecule detection, $J_0(\mathbf{x}) = \lambda c(\mathbf{x})$ |
| $\mathbf{s}(t)$ | vector-valued concentration signal, see Eq. (4) |
| $s(t)$ | $= \|\mathbf{s}\|$ train of molecular binding events |
| $J_0(\mathbf{x})$ | $= \lambda c(\mathbf{x})$ rate of molecule detection for a single receptor |
| $J(\mathbf{x})$ | $= \mathbb{E}(s \,\|\, \mathbf{x})$ rate of molecule detection, provided target at $\mathbf{x}^* = \mathbf{x}_0 + \mathbf{x}$ |
| | $\approx \lambda[c(\mathbf{x}_0) + (a^2/4)\nabla^2 c(\mathbf{x}_0)]$ |
| $\mathrm{SNR}^{\mathrm{SC}}$ | $= a^2 t \lambda \|\nabla c\|^2/c$ signal-to-noise ratio for spatial comparison, Eq. (1), |
| $\mathrm{SNR}^{\mathrm{TC}}$ | $= v^2 t^3 \lambda \|\nabla c\|^2/c$ signal-to-noise ratio for temporal comparison, Eq. (2), |
| $R_0$ | initial target distance |
| $R_{\mathrm{target}}$ | target size |
| $R_{\mathrm{max}}$ | radius of simulation domain (with absorbing boundary conditions) |
| $t_{\mathrm{turn}}$ | time till first turn for case $D_{\mathrm{rot}} = 0$ |
| $t_{\mathrm{ballistic}}$ | characteristic time for ballistic motion |
| $P_{\mathrm{reached}}$ | probability to reach target |
| $I_\varphi$ | directional information [in bits] |
| $I_\varphi^*$ | quasi-steady state of directional information, see Fig. 3 |
| $I_\varphi^{\mathrm{TC+SC}}$ | cumulated directional information: contribution due to TC and SC |
| $I_\varphi^{D_{\mathrm{rot}}}$ | cumulated directional information: contribution due to motility noise |
| $I_\varphi^{\mathrm{adv}}$ | cumulated directional information: contribution due to advection |
| $I_\varphi^{\mathrm{TC}}$ | cumulated directional information: contribution due to TC |
| $I_\varphi^{\mathrm{SC}}$ | cumulated directional information: contribution due to SC |
| $\mathbb{E}(s)$ | $= \langle J \rangle /2 \approx \lambda[\langle c \rangle + (a^2/4) \langle \nabla^2 c \rangle]$ expectation value of molecular binding events $s(t)$ |
| $\mathbb{E}(\mathbf{s})$ | $= \lambda a \langle \nabla c \rangle /2$ expectation value of the vector signal $\mathbf{s}(t)$ |
| $\mathbb{E}(\mathbf{s}\|s > 0)$ | $= \mathbb{E}\mathbf{s}\, s/\langle J \rangle = s\lambda a \langle \nabla c \rangle /(2 \langle J \rangle)$ conditional expectation value of vector signal $\mathbf{s}(t)$ |

TABLE S5. List of symbols.

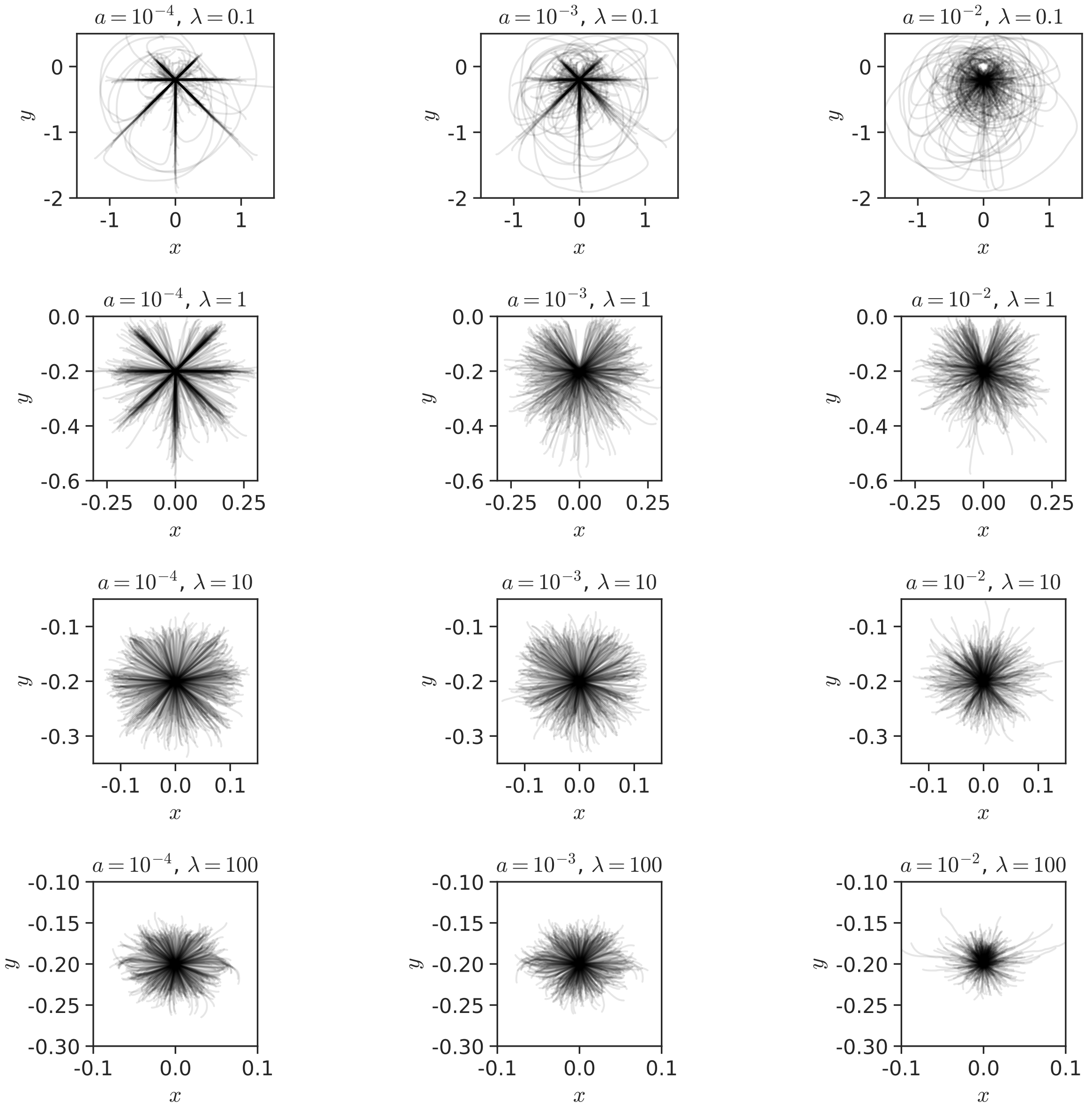


FIG. S1. **Stereotypic navigation in absence of motility noise: trajectories until first turn.** Overlay of trajectories of simulated infotaxis agents in the absence of motility noise $D_{\mathrm{rot}} = 0$ for different parameter values of agent size $a$ and rate constant $\lambda$ of molecule detection as analyzed in Fig. 2. Only initial parts of trajectories up to their first turning points are shown (black, transparent).

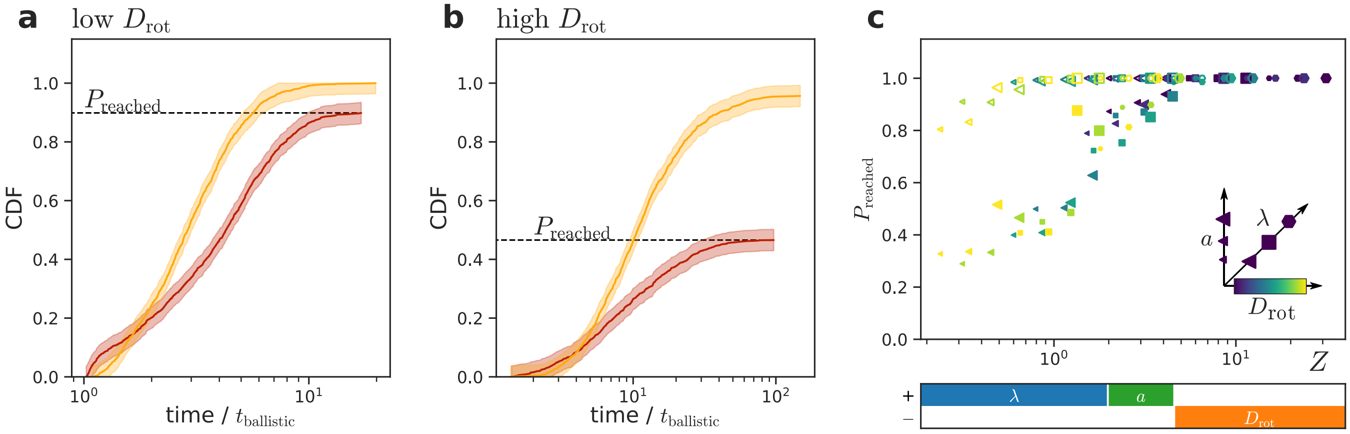


FIG. S2. **Probability to find the target with motility noise. a, b** Cumulative density function as function of normalized time $t/t_{\mathrm{ballistic}}$ with $t_{\mathrm{ballistic}} = R_{\mathrm{target}}/v$ for cases of low motility noise (panel a) and high motility noise (panel b) of the probability to find the target within time $t$, where $P_{\mathrm{reached}}$ is the probability for an agent to eventually find the target. (orange: maximum-likelihood strategy, red: infotaxis; 68% confidence interval based on DKW inequality [65]). **c** The probability $P_{\mathrm{reached}}$ to eventually find the target collapses on a power-law, $P_{\mathrm{reached}}(Z)$ with $Z = \lambda^{0.44}\,D_{\mathrm{rot}}^{-0.40}\,a^{0.16}$ (maximum-likelihood strategy: open symbols, infotaxis: filled symbols; agent parameters encoded according to coordinate system in the insert, see Fig. 3 for details). Power-law exponents determined for infotaxis by linear fit. Parameters (unless otherwise stated): a: $a = 0.01$, $\lambda = 1$, $D_{\mathrm{rot}} = 0.01$; b: same as panel a but $D_{\mathrm{rot}} = 0.5$; c: $a = 10^{-4} - 10^{-2}$, $\lambda = 1 - 100$, $D_{\mathrm{rot}} = 5 \cdot 10^{-3} - 1$.

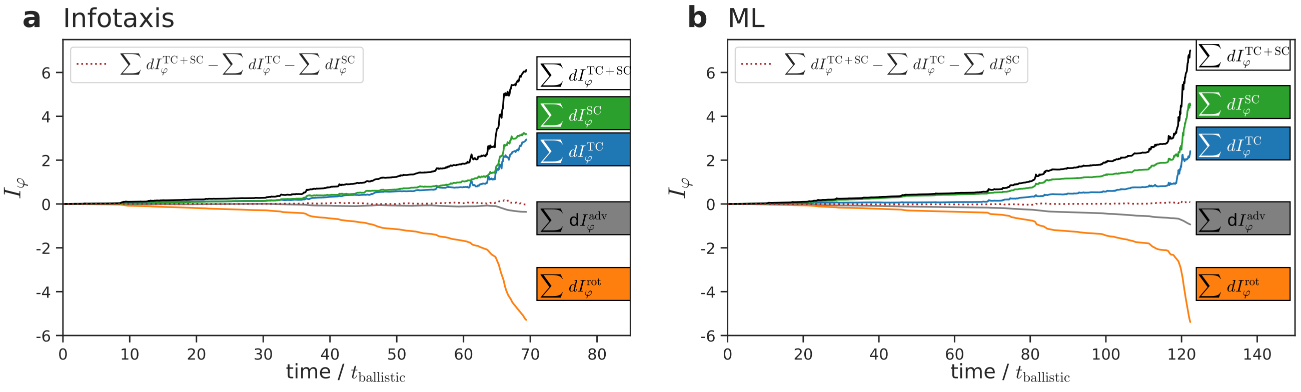


FIG. S3. **Information decomposition. a** Contributions to the cumulated change in directional information due to rotational diffusion $\sum \Delta I_\varphi^{\mathrm{rot}}$ (orange), advection $\sum \Delta I_\varphi^{\mathrm{adv}}$ (gray), concentration sensing (TC) $\sum \Delta I_\varphi^{\mathrm{TC}}$ (blue), gradient sensing (SC) $\sum \Delta I_\varphi^{\mathrm{SC}}$ (green), and full update step (TC+SC) $\sum \Delta I_\varphi^{\mathrm{TC+SC}}$ (white) as function of time $t$ for a typical simulation of an infotaxis agent. Information change contributions correspond to the different terms of the time-evolution equation for the likelihood, Eq. (5), and were computed according to Eqs. (S5)-(S9), see Section 'Information Decomposition'. The difference $\sum \Delta I_\varphi^{\mathrm{TC+SC}} - \Delta I_\varphi^{\mathrm{TC}} - \Delta I_\varphi^{\mathrm{SC}}$ (dashed) is approximately zero, confirming the validity of the information decomposition. **b** Same for a chemotactic agent using a maximum-likelihood decision strategy. Parameters: $a = 10^{-2}$, $\lambda = 1$, $D_{\mathrm{rot}} = 1$.

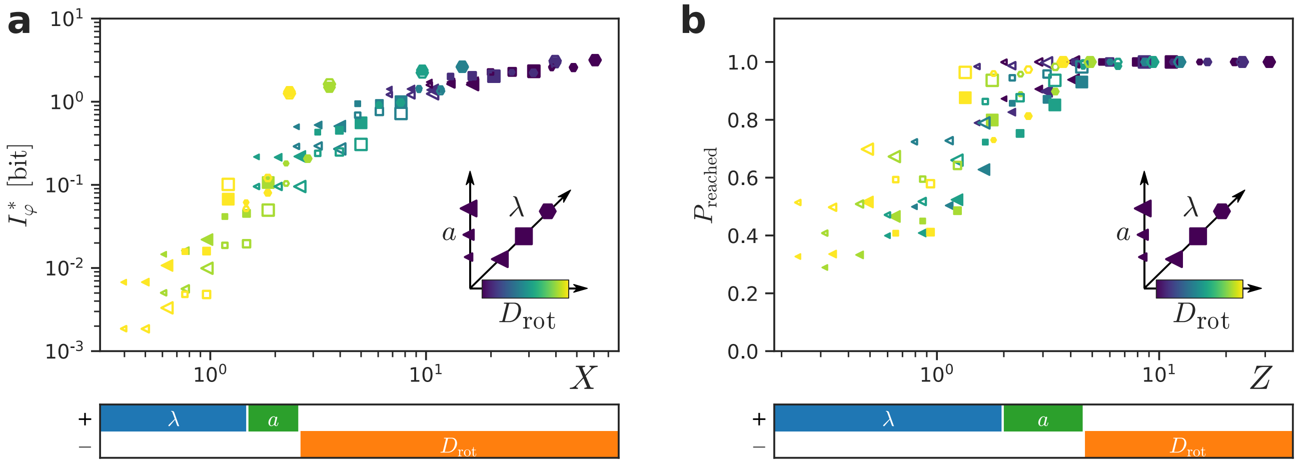


FIG. S4. **Similar power-laws for larger search domain.** Simulation results analogous to Fig. 3d and 3f, but for a larger search domain with $R_{\max} = 0.98$, reveal similar power-laws. Due to the long simulation time, the number of simulation can be as low as 38 for large $D_{\mathrm{rot}}$. **a** Quasi-steady states of directional information $I^*_\varphi(X)$ of infotaxis agents analogous to Fig. 3d but for $R_{\max} = 0.98$ (open symbols), and for comparison $R_{\max} = 0.48$ (filled symbols, same as Fig. 3d) using the same empirical power-law $X \sim \lambda^{0.28}\, D_{\mathrm{rot}}^{-0.62}\, a^{0.10}$ as determined for $R_{\max} = 0.48$ in Fig. 3d. Re-fitting the power-law with the data for $R_{\max} = 0.98$ gives similar exponents, $X \sim \lambda^{0.32}\, D_{\mathrm{rot}}^{-0.58}\, a^{0.10}$. **b** Probability to eventually find the target $P_{\mathrm{reached}}(Z)$ is larger for a search domain with $R_{\max} = 0.98$ (open symbols) as compared to a smaller search domain with $R_{\max} = 0.48$ (closed symbols, same as Fig. 3f), but collapses on the same power-law, $Z \sim \lambda^{0.44}\, D_{\mathrm{rot}}^{-0.40}\, a^{0.16}$ as determined for $R_{\max} = 0.48$ in Fig. 3f. Re-fitting the power-law for $R_{\max} = 0.98$ gave the same exponents. Parameters: as in Fig. 3, except $R_{\max}$ as stated.

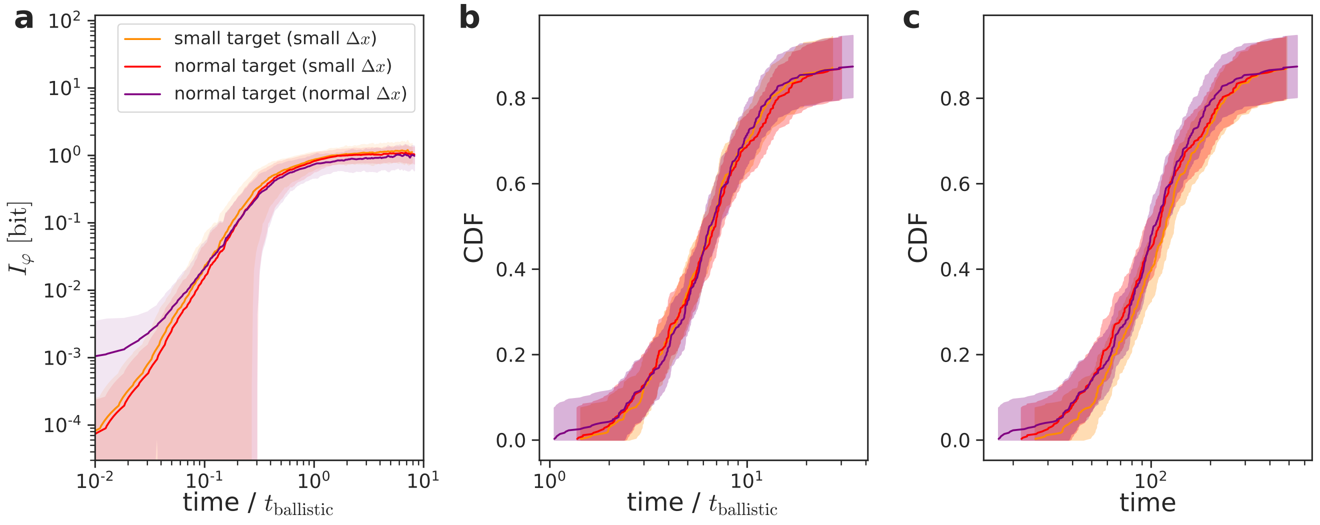


FIG. S5. **Similar results for smaller targets and smaller space discretization.** Directional information $I_\varphi(t)$ as function of search time (analogous to Fig. 3c), and cumulative density function of first-passage times (CDF) (analogous to Fig. 3e) for infotaxis agents, using two different sizes $R_{\mathrm{target}}$ of the target: $R_{\mathrm{target}} = 0.04$ (purple, red), and smaller target with $R_{\mathrm{target}} = 0.02$ (dark orange). To ensure numerical stability, implementation of a smaller target size required the use of a smaller space discretization $\Delta x$, which is also used for the red large target. **a** Directional information $I_\varphi(t)$ as function of search time for two target sizes. The difference in $I_\varphi(t)$ for the two target sizes at very short times is a result of the different space discretization used for the respective simulations. Note that the quasi-steady state $I^*_\varphi$ of directional information was at least five-times larger than this difference even for the most unfavorable parameters. **b** Cumulative density function of normalized first-passage time for two target sizes. There is almost no difference in the CDF for both target sizes for agents with short first-passage times, i.e. that move almost ballistically to the target. **c** Same as panel a, but without normalization of first-passage times. There is almost no difference in the CDF for both target sizes for agents with long first-passage times. Parameters: $a = 10^{-3}$, $D_{\mathrm{rot}} = 0.05$, $\lambda = 10$; normal target: $\Delta x = 0.01$, $R_{\mathrm{target}} = 0.04$; small target: $\Delta x = 0.005$, $R_{\mathrm{target}} = 0.02$.

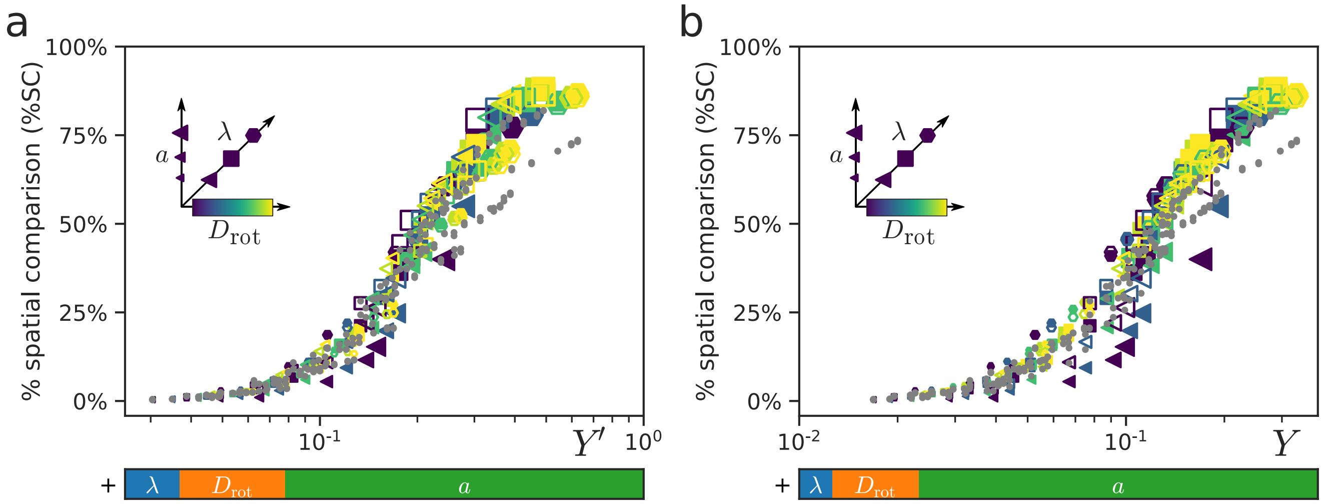


FIG. S6. **Similar information decomposition for relative importance of spatial comparison for exponential concentration field.** Relative importance of spatial comparison %SC, computed by metering information flow as in Fig. 4a in the main text, yet for agents searching in an exponential concentration field as given by Eq. (S18) (instead of the usual concentration field $c(\mathbf{x}) = 1/|\mathbf{x}|$), plotted as function of an effective master parameter. Filled symbols correspond to infotaxis agents, open symbols to maximum-likelihood agents, agent parameters encoded according to inset; gray dots correspond to original data from Fig. 4a, here re-shown for reference. **a** Relative importance $\%\mathrm{SC}(Y')$ plotted as function of a new master parameter $Y' = \lambda^{0.11} a^{0.69} D_{\mathrm{rot}}^{0.20}$, with exponents obtained by a fit to the simulation data for infotaxis agents searching in an exponential concentration field, see also Table S2. **b** Same data as in panel **a**, but plotted as function of the original master parameter $Y$ from Fig. 4a in the main text. Parameters: $a = 10^{-3} - 2 \cdot 10^{-2}$, $\lambda = 1 - 100$, $D_{\mathrm{rot}} = 0.1 - 1$.

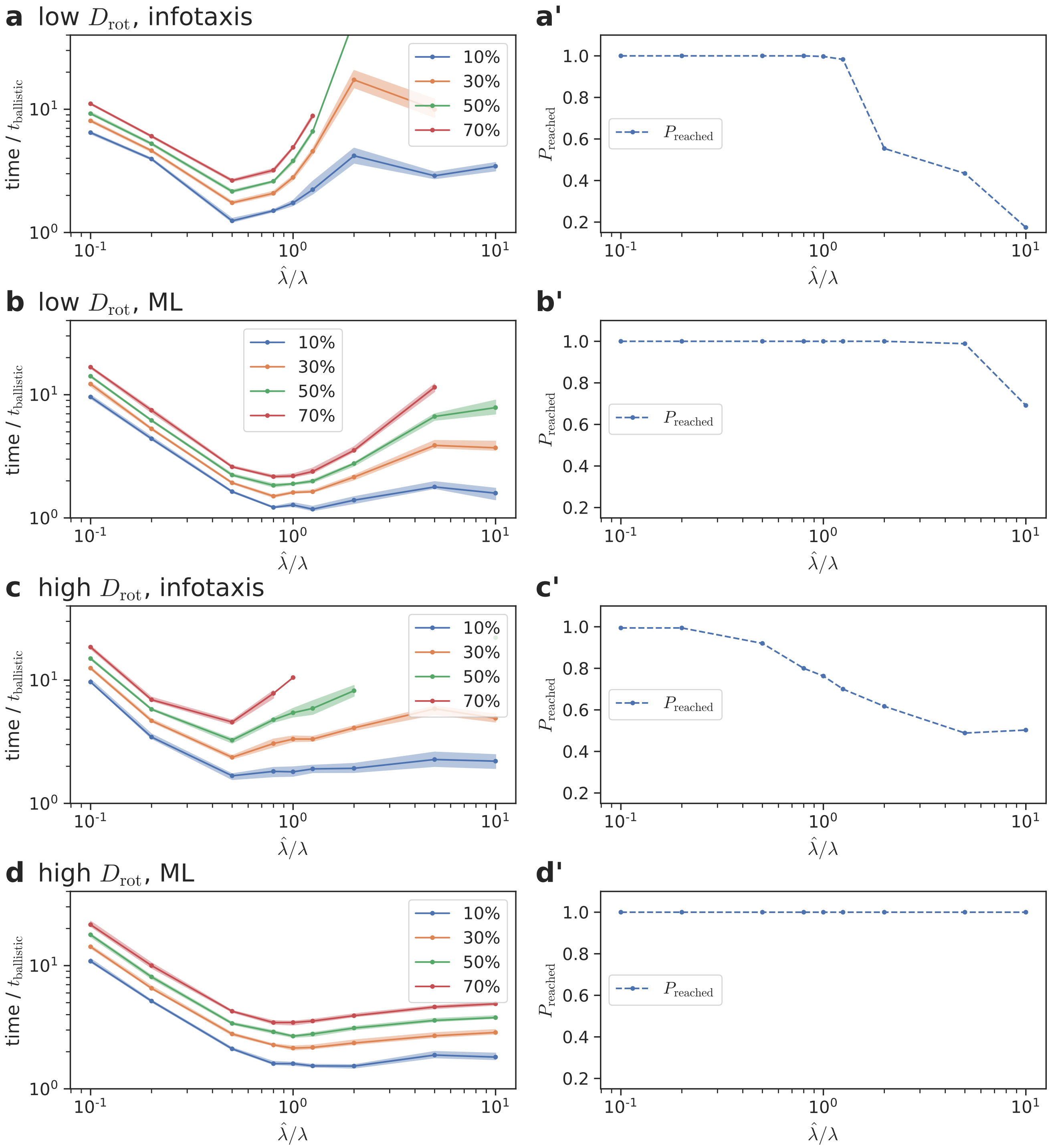


FIG. S7. **Search performance for inaccurate model of environment: rate constant of molecule detection.** Performance of chemotactic agents that assume an inaccurate value $\widehat{\lambda}$ of the rate constant of molecule detection in their update step, quantified in terms of quantiles of the first-passage time $t$ (10%: blue, 30%: orange, 50%: green, 70%: red; left axis), and the probability $P_{\text{reached}}$ to eventually reach the target (primed panels) for different scenarios. **a** Infotaxis agents with low motility noise $D_{\text{rot}} = 0.01$. **b** Maximum-likelihood agents with low motility noise $D_{\text{rot}} = 0.01$. **c** Infotaxis agents with high motility noise $D_{\text{rot}} = 0.01$. **d** Maximum-likelihood agents with high motility noise $D_{\text{rot}} = 0.1$. Parameters: $a = 10^{-3}$, $\lambda = 10$, a, b: $D_{\text{rot}} = 0.01$, c, d: $D_{\text{rot}} = 0.1$.

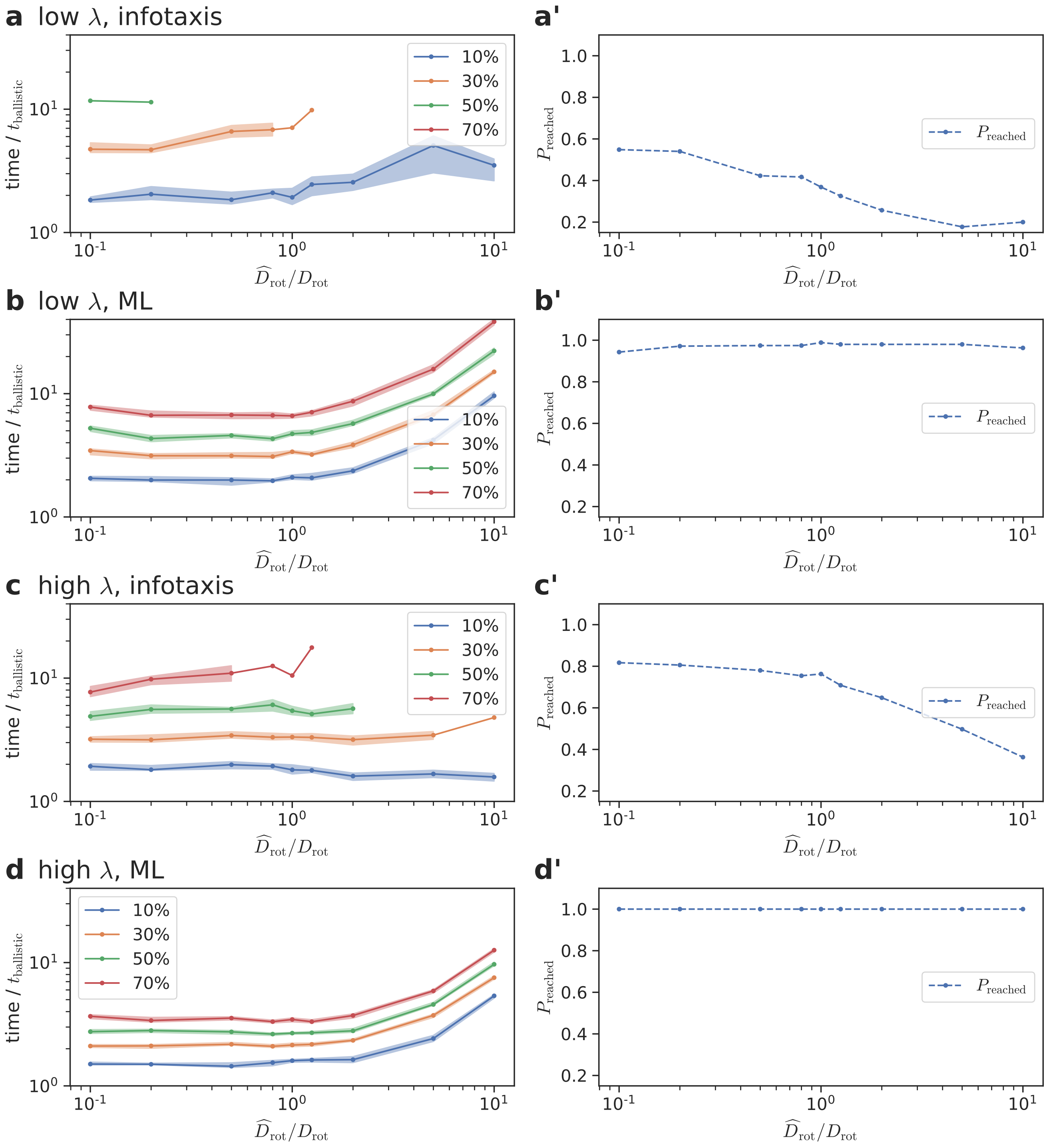


FIG. S8. **Search performance for inaccurate model of environment: rotational diffusion coefficient.** Performance of chemotactic agents that assume an inaccurate value $\widehat{D}_{\mathrm{rot}}$ of their rotational diffusion coefficient different from the true value $D_{\mathrm{rot}}$ in their prediction step, quantified in terms of quantiles of the first-passage time $t$ (10%: blue, 30%: orange, 50%: green, 70%: red; left axis), and the probability $P_{\mathrm{reached}}$ to eventually reach the target (primed panels) for different scenarios analogous to Fig. S7. **a** Infotaxis agents with low sensing noise $\lambda = 1$. **b** Maximum-likelihood agents with low sensing noise $\lambda = 1$. **c** Infotaxis agents with high sensing noise $\lambda = 10$. **d** Maximum-likelihood agents with high sensing noise $\lambda = 10$. Parameters: $a = 10^{-3}$, $D_{\mathrm{rot}} = 0.1$, a, b: $\lambda = 10$, c, d: $\lambda = 1$.

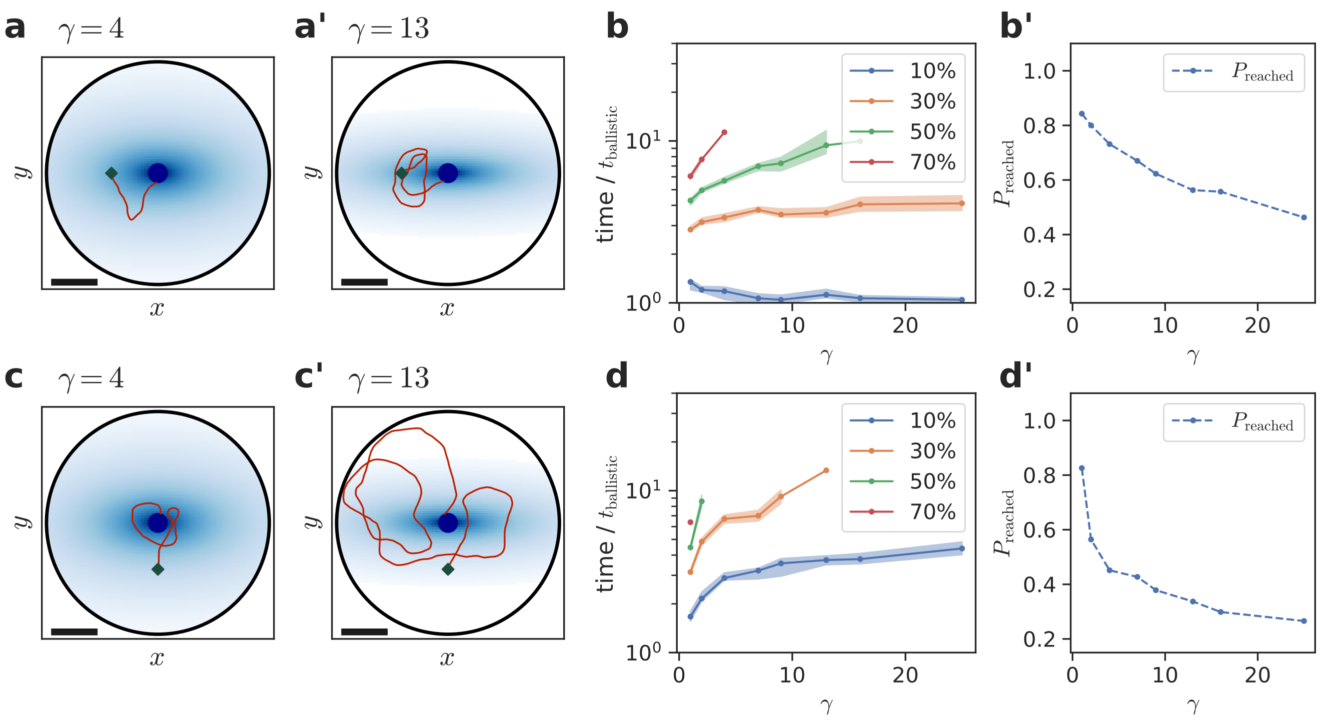


FIG. S9. **Search performance for inaccurate model of environment: non-radial concentration field.** Performance of chemotactic agents that wrongly assume a radially symmetric concentration field ($c(\mathbf{x}) = 1/|\mathbf{x}|$), while the true concentration field is compressed along the $y$-direction by a factor $\gamma^{1/2}$ (given by $c(x, y) = 1/\sqrt{x^2 + \gamma\, y^2}$). **a** Visualization of concentration fields (shades of blue), with typical infotaxis trajectory (black) and initial position $\mathbf{x}(t = 0) = (-R_0, 0)^T$ (diamond) for two values of $\gamma$: $\gamma = 4$ (**a**), and $\gamma = 13$ (**a'**). **b** Search performance is quantified in terms of quantiles of the first-passage time $t$ (10%: blue, 30%: orange, 50%: green, 70%: red; left axis), and the probability $P_{\text{reached}}$ to eventually reach the target. **c** Analogous to panel **a**, but for different initial position $\mathbf{x}(t = 0) = (0, -R_0)^T$. **d** Analogous to panel **b**, but for different initial position $\mathbf{x}(t = 0) = (0, -R_0)^T$. Parameters: $a = 10^{-3}$, $D_{\text{rot}} = 0.01$, $\lambda = 1$.

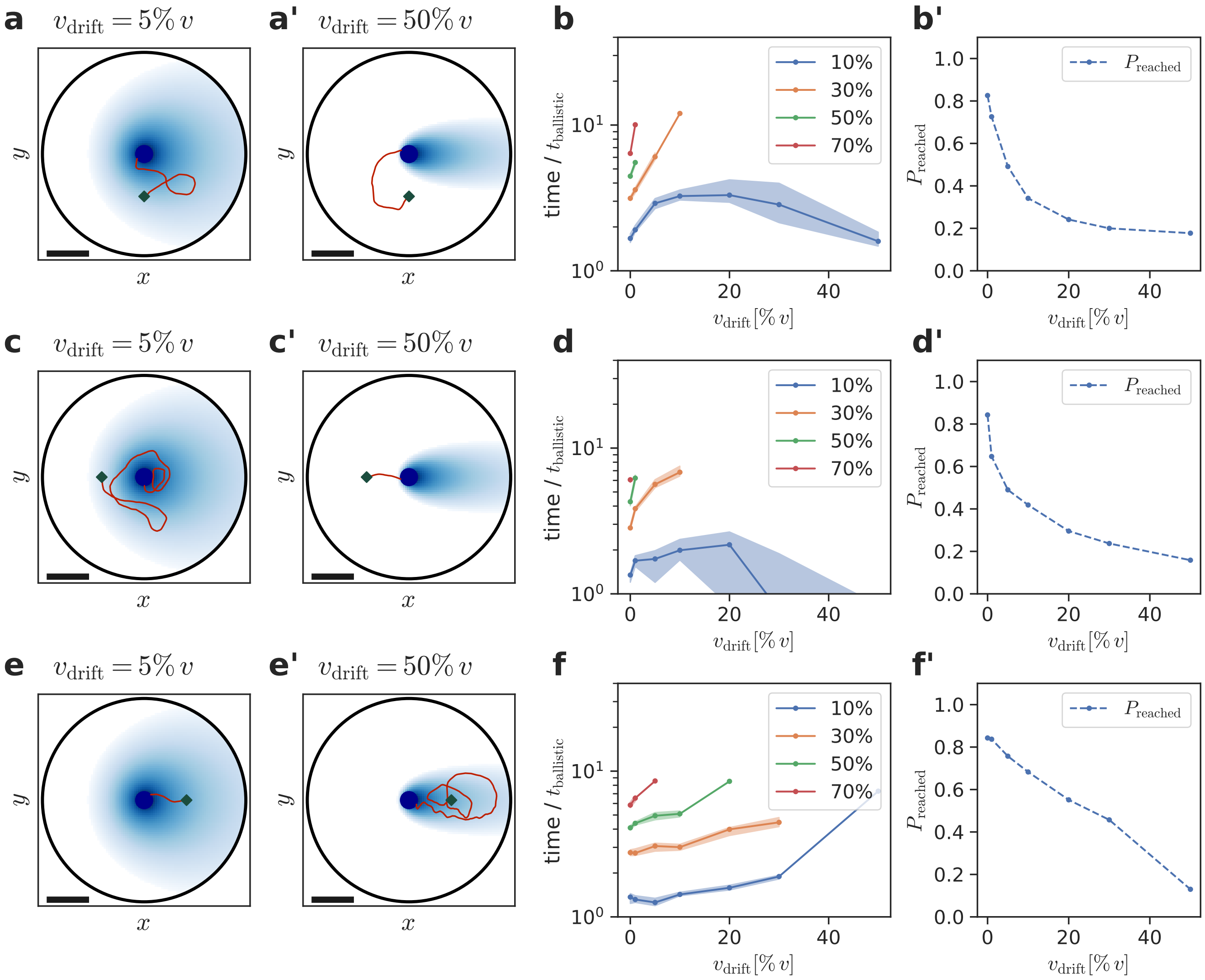


FIG. S10. **Search performance for inaccurate model of environment: concentration field distorted by drift.** Performance of chemotactic agents that wrongly assume a radially symmetric concentration field ($c(\mathbf{x}) = 1/|\mathbf{x}|$), while the true concentration field is distorted by drift in $x$-direction with drift speed $v_{\rm drift}$ (given by $c(x,y) = \exp\left[v_{\rm drift}/(2\,D_0)\,(x - |\mathbf{x}|)\right]/|\mathbf{x}|$). **a** Visualization of concentration fields (shades of blue), with typical infotaxis trajectory (black) and initial position $\mathbf{x}(t=0) = (0, -R_0)^T$ (diamond) for two values of the drift speed $v_{\rm drift}$: $v_{\rm drift} = 0.05\, v_0$ (**a**), and $v_{\rm drift} = 0.5\, v_0$ (**a'**). **b** Search performance is quantified in terms of quantiles of the first-passage time $t$ (10%: blue, 30%: orange, 50%: green, 70%: red; left axis), and the probability $P_{\rm reached}$ to eventually reach the target (primed panels). **c, e** Analogous to panel **a**, but for different initial position $\mathbf{x}(t=0) = (-R_0, 0)^T$ (**c**), and $\mathbf{x}(t=0) = (R_0, 0)^T$ (**e**). **d** Analogous to panel **b**, but for different initial position $\mathbf{x}(t=0) = (-R_0, 0)^T$ (**d**), and $\mathbf{x}(t=0) = (R_0, 0)^T$ (**f**). Parameters: $a = 10^{-3}$, $D_{\rm rot} = 0.01$, $\lambda = 1$, $D_0 = 10^{-4}$.

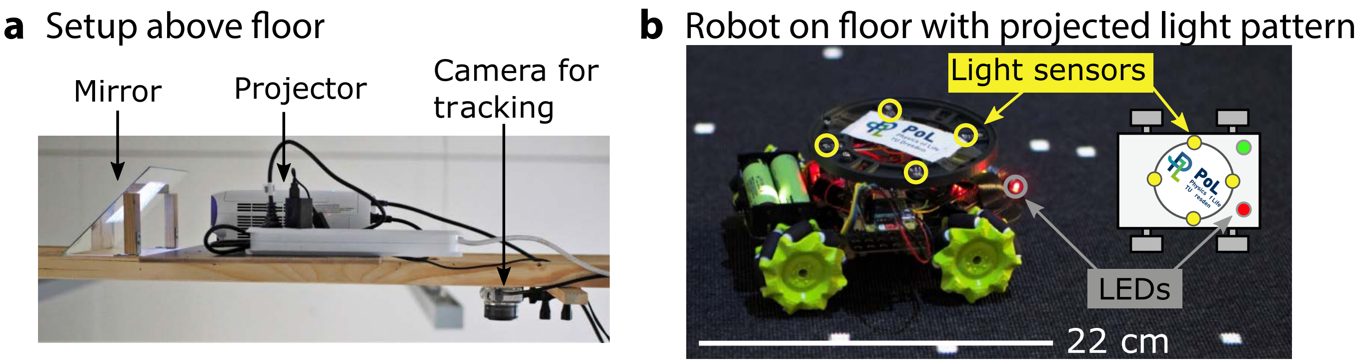


FIG. S11. **Experimental setup.** **a** A beamer mounted on a stand projects a pixel pattern on the search arena on the floor via a diagonally aligned mirror. A camera on the same stand tracks red and green LEDs on the robot to determine its the position and orientation. **b** The robot is equipped with a circular array of 4 light sensors, which feed into a Raspberry Pi micro-computer. A red and green LED on the robot serve as tracking aids.

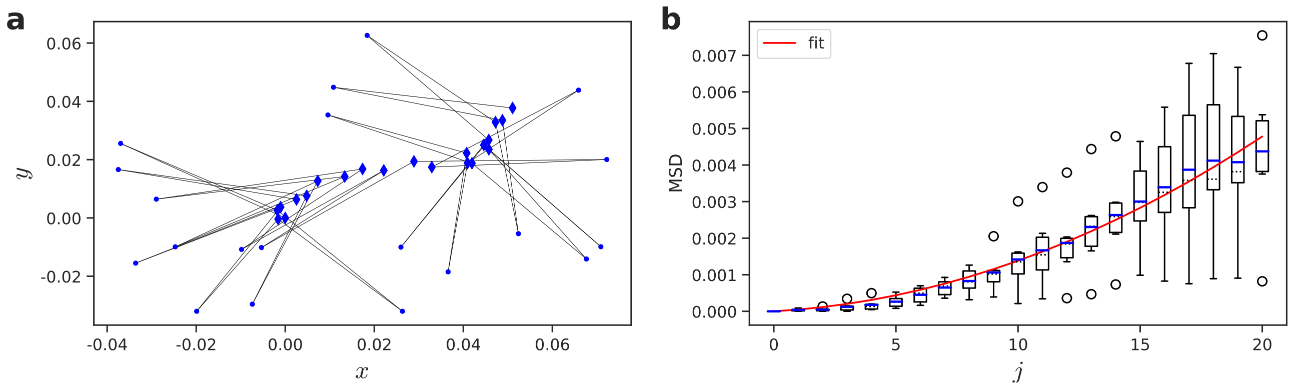


FIG. S12. **Measuring motility noise of the robot.** **a** Diamonds show the positions $\mathbf{x}_j$ of the robot after the $j$th back-and-forth maneuver, while dots depict the intermediate positions at which the robot reverses its direction. The thin black line indicates the movement of the robot. **b** Mean square displacement (MSD) $\langle|\mathbf{x}_j - \mathbf{x}_0|^2\rangle$ for the motility task shown in panel **a** as function of $j$ (dashed line: median; blue line: mean; box: 1/4 and 3/4 quantiles; whiskers: 1.5 IQR; points: outliers; $n = 6$). The red line shows a least-square fit $\mathrm{MSD}(j) \approx c\,j + d\,j^2$, from which we estimate the parameters $c = (4 \pm 7) \cdot 10^{-5}$ and $d = (1.0 \pm 0.5) \cdot 10^{-5}$ (errors equal one standard deviation calculated from Hessian matrix).

### H. SUPPLEMENTARY INFORMATION

**Supplementary Materials**
Supplementary Figs. S1-S12, including extended numerical and analytical methods.

**Supplementary videos**
Supplementary videos show the time evolution of the likelihood map $p(x, y, t)$ of target position in the laboratory frame of reference for simulated agents and robot experiments (agent: green disk drawn to scale, or green cross if too small; target: blue). Panels to the right show the directional information $I_\varphi(t)$ as function of time $t$, where the red line indicates current time, as well as a cartoon representation of the agent visualizing molecular binding events relative to the material frame of the agent. The coordinate system inside the agent depicts the laboratory frame of reference.

**Supplementary video 1**
Agent without motility noise performing infotaxis. Parameters: $a = 10^{-4}$, $\lambda = 1$, $D_{\mathrm{rot}} = 0$.

**Supplementary video 2**
Agent without motility noise using maximum-likelihood strategy. Parameters: $a = 10^{-4}$, $\lambda = 1$, $D_{\mathrm{rot}} = 0$.

**Supplementary video 3**
Agent with motility noise performing infotaxis. Parameters: $a = 10^{-2}$, $\lambda = 1$, $D_{\mathrm{rot}} = 0.1$.

**Supplementary video 4**
Agent with motility noise using maximum-likelihood strategy. Parameters: $a = 10^{-2}$, $\lambda = 1$, $D_{\mathrm{rot}} = 0.1$.

**Supplementary video 5**
Robot experiment for maximum-likelihood strategy.

*


[1] D. B. Dusenbery, Biophys. J. **74**, 2272 (1998).
[2] K. Y. Wan and G. Jékely, Phil. Trans. R. Soc. B **376**, 20190758 (2021).
[3] L. Alvarez, B. M. Friedrich, G. Gompper, and U. B. Kaupp, Trends Cell Biol. **24**, 198 (2014).
[4] S. H. Zigmond, Nature **249**, 450 (1974).
[5] C. A. Parent and P. N. Devreotes, Science **284**, 765 (1999).
[6] P. J. Van Haastert and M. Postma, Biophys. J. **93**, 1787 (2007).
[7] G. Amselem, M. Theves, A. Bae, C. Beta, and E. Bodenschatz, Phys. Rev. Lett. **109**, 1 (2012).
[8] A. Nakajima, S. Ishihara, D. Imoto, and S. Sawai, Nat. Commun. **5**, 5367 (2014).
[9] H. C. Berg and D. A. Brown, Nature **239**, 500 (1972).
[10] R. M. Macnab and D. E. Koshland, Proc. Natl. Acad. Sci. U.S.A. **69**, 2509 (1972).
[11] J. E. Segall, S. M. Block, and H. C. Berg, Proc. Nat. Acad. Sci. U.S.A. **83**, 8987 (1986).
[12] D. R. Brumley, F. Carrara, A. M. Hein, Y. Yawata, S. A. Levin, and R. Stocker, Proc. Natl. Acad. Sci. U.S.A. **166**, 10792 (2019).
[13] H. H. Mattingly, K. Kamino, B. B. Machta, and T. Emonet, Nat. Phys. **17**, 1426 (2021).
[14] J. F. Jikeli, L. Alvarez, B. M. Friedrich, L. G. Wilson, R. Pascal, R. Colin, M. Pichlo, A. Rennhack, C. Brenker, and U. B. Kaupp, Nat. Commun. **6**, 7985 (2015).
[15] H. C. Berg and E. M. Purcell, Biophys. J. **20**, 193 (1977).
[16] R. G. Endres and N. S. Wingreen, Proc. Natl. Acad. Sci. U. S. A **105**, 15749 (2008).
[17] B. Hu, W. Chen, W.-J. Rappel, and H. Levine, Phys. Rev. Lett. **105**, 048104 (2010).
[18] W. Bialek and S. Setayeshgar, Proc. Natl. Acad. Sci. U.S.A. **102**, 10040 (2005).
[19] K. Kaizu, W. De Ronde, J. Paijmans, K. Takahashi, F. Tostevin, and P. R. ten Wolde, Biophys. J. **106**, 976 (2014).
[20] T. Mora and N. S. Wingreen, Phys. Rev. Lett. **104**, 248101 (2010).
[21] A. M. Hein, D. R. Brumley, F. Carrara, R. Stocker, and S. A. Levin, J. R. Soc. Interface **13** (2016).
[22] D. Fuller, W. Chen, M. Adler, A. Groisman, H. Levine, W.-J. Rappel, and W. F. Loomis, Proc. Natl. Acad. Sci. U.S.A. **107**, 9656 (2010).
[23] K. L. Thornton, J. K. Butler, S. J. Davis, B. K. Baxter, and L. G. Wilson, Nat. Commun. **11**, 4453 (2020).
[24] K. Nakamura and T. J. Kobayashi, Phys. Rev. Res. **4**, 013120 (2022).
[25] M. Vergassola, E. Villermaux, and B. Shraiman, Nature **445**, 406 (2007).
[26] A. Celani and M. Vergassola, Proc. Natl. Acad. Sci. U.S.A. **107**, 1391 (2010).
[27] C. Barbieri, S. Cocco, and R. Monasson, EPL **94**, 20005 (2011).
[28] B. Hartl, M. Hübl, G. Kahl, and A. Zöttl, Proc. Natl. Acad. Sci. U.S.A **118**, e2019683118 (2021).
[29] N. Rigolli, N. Magnoli, L. Rosasco, and A. Seminara, eLife **11**, e72196 (2022).
[30] G. Reddy, V. N. Murthy, and M. Vergassola, Annu. Rev. Condens. Matter Phys. **13**, 191 (2022).
[31] A. Loisy and C. Eloy, Proc. R. Soc. A **478**, 20220118 (2022).
[32] A. Loisy and R. A. Heinonen, The European Physical Journal E **46**, 17 (2023).
[33] A. R. Cassandra, L. P. Kaelbling, and J. A. Kurien, in *Proceedings of IEEE/RSJ International Conference on Intelligent Robots and Systems. IROS'96* (IEEE, 1996), vol. 2, pp. 963–972.
[34] A. Auconi, M. Novak, and B. M. Friedrich, EPL **138**, 12001 (2022).
[35] A. Bain and D. Crisan, *Fundamentals of stochastic filtering* (Springer, 2009).
[36] J. Kromer, S. Märcker, S. Lange, C. Baier, and B. M. Friedrich, PLoS Comp. Biol. **14**, 1 (2018).
[37] S. P. Strong, B. Freedman, W. Bialek, and R. Koberle, Phys. Rev. E **57**, 4604 (1998).
[38] T. Mora and I. Nemenman, Phys. Rev. Lett. **123**, 198101 (2019).
[39] M. Novak and B. M. Friedrich, New J. Phys. **23**, 043026 (2021).
[40] R. O. P. Ramakrishnan and B. M. Friedrich, EPL (2023).
[41] J. H. Wheeler, K. R. Foster, and W. M. Durham, bioRxiv pp. 2024–02 (2024).
[42] R. Thar and M. Kühl, Proc. Natl. Acad. Sci. U.S.A. **100**, 5748–5753 (2003).
[43] A. Einstein, Ann. Phys. **517**, 248 (1906/2005).
[44] H. Crenshaw, Am. Zool. **36**, 608 (1996).
[45] B. M. Friedrich and F. Jülicher, Proc. Natl. Acad. Sci. U.S.A. **104**, 13256 (2007).
[46] R. Z. Tan and K.-H. Chiam, PLoS Comput. Biol. **14**, e1005966 (2018).
[47] Moraud, Front. Neurorobot. (2010).
[48] J.-B. Masson, Proc. Natl. Acad. Sci. U.S.A **110**, 11261 (2013).
[49] N. Voges, A. Chaffiol, P. Lucas, and D. Martinez, PLoS Comput. Biol. **10**, e1003861 (2014).
[50] U. Dobramysl and D. Holcman, Phys. Rev. Lett. **125**, 148102 (2020).
[51] K. Nakamura and T. J. Kobayashi, *Gradient sensing limit of a cell when controlling the elongating direction* (2024), 2405.04810.
[52] A. Alonso and J. B. Kirkegaard, arXiv preprint arXiv:2310.10531v2 (2024).
[53] R. Stocker, Science **338**, 628 (2012).
[54] S. Lange and B. M. Friedrich, PLoS Comput. Biol. **17**, e1008826 (2021).
[55] J. D. Rodríguez, D. Gómez-Ullate, and C. Mejía-Monasterio, Euro. Phys. J. Spec. Topics **226**, 2407 (2017).

[56] A. Celani, E. Villermaux, and M. Vergassola, Phys. Rev. X **4**, 041015 (2014).
[57] K. V. B. Verano, E. Panizon, and A. Celani, Proceedings of the National Academy of Sciences **120**, e2304230120 (2023).
[58] P. Romanczuk, M. Bär, W. Ebeling, B. Lindner, and L. Schimansky-Geier, Eur. Phys. J. Spec. Top. **202**, 1 (2012).
[59] A. M. Hein and S. A. McKinley, Proc. Natl. Acad. Sci. U.S.A. **109**, 12070 (2012).
[60] J. A. Kromer, A. Auconi, and B. M. Friedrich, ChemNanoMat **7**, 1057 (2021).
[61] B. M. Friedrich, Phys. Biol. **5**, 026007 (2008).
[62] K. J. Astrom et al., Journal of mathematical analysis and applications **10**, 174 (1965).
[63] R. D. Smallwood and E. J. Sondik, Operations research **21**, 1071 (1973).
[64] C. Baier and J.-P. Katoen, *Principles of model checking* (MIT press, 2008).
[65] P. Massart, Ann. Probab. **18** (1990).
